# Cyst-like intraluminal hyaluronic acid accumulation inhibits breast epithelial cell invasion via pressure-driven ECM compaction

**DOI:** 10.64898/2026.11.25.769205

**Authors:** B.E. Schutrum, K. Kim, D.A. Rivera, J. Kim, S. Balakrishnan, S. Joshi, A.A. Shimpi, S. Girardo, V. Mango, N. Bouklas, S. Adie, C.B. Schaffer, J.M. Schwarz, C. Fischbach

**Affiliations:** Nancy E. and Peter C. Meinig School of Biomedical Engineering, Cornell University, Ithaca, NY, USA; Department of Mathematics, University of British Columbia, Vancouver, BC, Canada; Sibley School of Mechanical and Aerospace Engineering, Cornell University, Ithaca, NY, USA; Max Planck Institute for the Science of Light, Erlangen, Germany; Department of Radiology, Memorial Sloan Kettering Cancer Center, New York, NY, USA; Indian Creek Farm, Ithaca, NY, USA; Physics Department, Syracuse University, Syracuse, NY, USA

## Abstract

While elevated hyaluronic acid (HA) biosynthesis by breast epithelial cells is implicated in tumorigenesis, HA can also contribute to benign tissue remodeling, including mammary morphogenesis and lumen formation in embryogenesis. What determines whether HA drives malignant invasion or benign remodeling remains unclear. To investigate, we embedded spheroids of HA synthase (HAS)-overexpressing breast epithelial cells in collagen hydrogels modeling dense breast stroma. Confocal and optical coherence microscopy showed that HAS overexpression produced different outcomes in single cells and spheroids: individual cells invaded into the collagen, whereas spheroids accumulated HA internally, causing osmotic swelling and hydrostatic pressure-driven lumen formation. An *in silico* cell-vertex model identified paracellular fluid flow as a mechanism of lumen formation, coinciding with confinement-induced reorganization of cell-cell adhesion strengths. *In situ* pressure sensors, confocal reflectance imaging, and finite element analysis showed that spheroid expansion compressed the surrounding collagen, inducing densification and a material phase transition that inhibited invasion. Co-culture and direct contact with matrix-remodeling stromal cells attenuated this effect and restored invasive potential. Although HA overproduction in single cells has been linked to malignancy, our findings identify a biophysical mechanism by which multicellular HA accumulation suppresses tumor invasion and show how stromal remodeling can override this protection. These results offer a potential explanation for the context-dependent relationship between fibrocystic breast changes and breast cancer risk.

## Introduction

Hyaluronic acid (HA) is a linear, anionic glycosaminoglycan that regulates physiological and pathological processes, including joint function and mammary tumorigenesis, through both its biochemical and biophysical properties^1–3^. HA is synthesized at the plasma membrane by three isoforms of the transmembrane enzyme hyaluronan synthase (HAS1-3), which differ in their regulation, catalytic rate and in the molecular weight (MW) of the HA they produce^4,5^. In breast tissue, HAS2 is the predominant isoform, and HAS2 and HAS3 synthesize HA at similar, higher rates than HAS1^4–7^. Epithelial overproduction of HA is associated with malignant progression, enhancing stemness, invasion, drug resistance, immune evasion, epithelial-to-mesenchymal transition, and metabolic alterations^8–11^. Most prior work linking HA to these phenotypes has focused on its biochemical effects, with signaling outcomes that depend on HA MW, and the consequent activation of distinct receptor-mediated signaling pathways involved in proliferation, survival, and motility^12^. However, HA is also a key structural component of the extracellular matrix whose material properties regulate cell fate and tissue function^3,13^, yet how these properties change within the breast tissue remains poorly understood.

HA’s stiff, extended polymer chains carry a dense fixed negative charge that attracts mobile counterions (e.g., Na⁺), which in turn draw water into the HA-rich region, making HA highly osmotic and capable of generating swelling and hydrostatic pressure^14,15^. During development, HA-dependent hydrostatic pressure directs cardiac valve formation^16^ and inner ear morphogenesis by driving lumen formation^13,17^. HA’s osmotic properties also shape adult tissue architecture, as HA has been implicated in the formation of ovarian carcinoma cysts^18,19^ and may similarly drive cyst formation in fibrocystic breast disease^20,21^, where breast cyst fluid is enriched in osmotically active proteoglycans^22,23^. Although fibrocystic disease is common (affecting over 50% of women) and can be painful^24,25^, it is benign and not associated with increased breast cancer risk. Thus, while the biochemical signaling pathways activated by HA can promote malignant behavior in individual epithelial cells, its biophysical properties may have opposing effects by driving morphogenic processes such as lumen formation and cyst development, which are not linked to malignancy. How microenvironmental conditions determine which of these outcomes dominates remains unclear.

To address this knowledge gap, *in vitro* models are needed that capture both multicellular organization and a physiologically relevant matrix, yet most existing models recapitulate neither. The effects of HA overproduction by breast epithelial cells have mainly been studied in 2D culture or with individual cells suspended in 3D matrices, which cannot reproduce multicellular biophysical processes such as pressure-driven lumen formation. Furthermore, although collagen-rich, fibrotic extracellular matrix (ECM) is a hallmark of both breast tumorigenesis^26,27^ and fibrocystic disease^20,25^, 3D studies of mammary lumen formation are often performed in reconstituted basement membrane (BM) such as Matrigel®, which lacks the biochemical, mechanical, and microstructural features of fibrillar ECM. BM systems cannot, for example, recapitulate the architecture and orientation of collagen fibers at the tumor boundary, which influence tumor invasiveness *in vivo*^28,29^. Engineered collagen hydrogels overcome these limitations by enabling precise control of bulk material properties^30–32^, cell-scale microarchitecture ^33,34^ and cell-mediated matrix remodeling^35,36^, so that the role of each parameter can be isolated. Combined with finite element analysis (FEA) of collagen networks^37,38^, which predicts local matrix deformations that regulate cell behavior, these hydrogels can reveal how matrix mechanics shape cell responses of interest including confinement-induced solid stress, which can induce collective invasion of tumor cells into the surrounding stroma^39,40^.

Here, we combined microfabricated collagen hydrogels, spheroid formation techniques, and genetic modulation of HA production by epithelial cells^8,41^ to mimic hyperplastic breast lesions. We used these mimics to examine the contrasting outcomes of non-malignant cyst formation and malignant invasion as a function of HA production. We further applied *in situ* measurements of hydrostatic pressure, laser ablation to disrupt the integrity of lumen-like structures, and computational modeling of lumen formation and matrix deformation to characterize how the biophysical properties of HA regulate breast epithelial behavior beyond its traditional role as a signaling molecule. By showing that HA physically remodels the surrounding matrix and thereby determines whether breast epithelial cells form benign lumens or invade as single cells, our findings underscore the importance of tissue context in HA-driven breast pathology and may help explain why HA-targeting therapies^42^ have shown limited success in breast cancer.

## Results

### Overexpression of HAS prevents 3D spheroid invasion while enabling spheroid expansion and internal lumen formation

To assess the effects of HA overproduction on breast epithelial cell phenotype from hyperplastic breast lesions, we generated spheroids from MCF10A cells with doxycycline-inducible HAS2 overexpression ^8^ (**Fig. 1a; Extended Data Fig. 1**). First, we confirmed that HAS2 overexpression promotes HA biosynthesis as well as malignant transformation in conventional 2D culture models by increasing mesenchymal cell morphologies with abundant filopodia and increased transwell invasion **(Extended Data Fig. 1)** consistent with previous results from our group and others ^8,10,43^. To test how HAS2 overexpression alters MCF10A behavior in 3D culture, more closely reflecting conditions *in vivo*, we prepared parental (control) and HAS2 spheroids using the hanging drop method and embedded them into microfabricated fibrillar 2.5 or 6 mg/mL collagen type I hydrogels (**Fig. 1b, Extended Data Fig. 2a-c**) to model pre-malignant and pathological ECM stiffness in the breast, respectively^32^. In contrast to collagen-embedded single cells^8^, HAS2 overexpression did not promote MCF10A spheroid invasion regardless of collagen concentration (**Fig. 1c**). Interestingly, HAS2 spheroids were significantly larger and contained translucent portions relative to parental controls in both collagen concentrations with a much more pronounced effect in 2.5 versus 6 mg/mL collagen. Because 6 mg/mL collagen type I more closely recapitulates the ECM density of fibrotic stroma^32^, we chose this condition to further elucidate the mechanisms leading to spheroid expansion. Analysis of DNA content and confocal micrographs demonstrated that HAS2 spheroid expansion was not due to a change in proliferation induced by the transduction protocol but caused by HA overproduction (**Extended Data Fig. 2d-e**). Spheroids expressing the HAS2 plasmid without doxycycline induction had significantly less expansion (<10 % vs.∼60% change in area) and were lacking translucent regions (**Extended Data Fig. 2f**). We observed the same effects with overexpression of HAS3 (**Extended Data Fig. 3a-e**), another HAS isoform with similar enzymatic activity to HAS2^4–6^. HAS2 spheroids embedded into reconstituted basement membrane also expanded with the characteristic translucent regions but more irregular spheroid morphologies (**Extended Data Fig. 4**). Given that spheroids expanded more in 2.5 vs. 6 mg/mL collagen and had less regular morphologies in reconstituted basement membrane, which is softer and less strain-stiffening than collagen, and because increased matrix density can lead to volumetric confinement^44^, these data suggest that HA overproduction leads to spheroid expansion which may be limited under conditions of increased ECM density. Taken together, our findings suggest that overexpression of HAS2 in multicellular, ECM-embedded models does not promote single-cell or collective invasion, a finding that contrasts previous studies with collagen-embedded single cells in which HAS2 overexpression promoted their invasion^8^.

**Figure 1:**
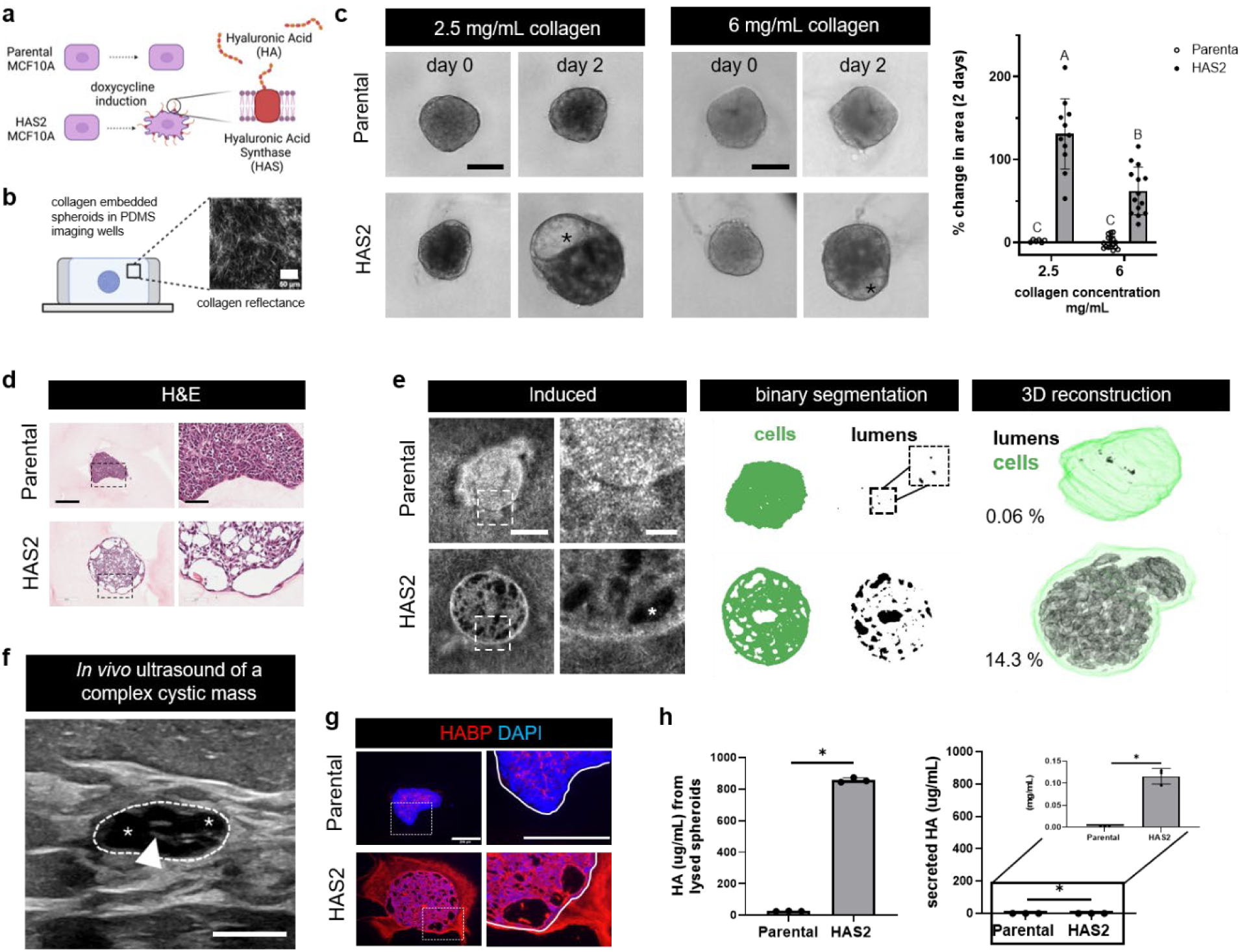
HAS2 overexpression promotes intraspheroidal lumen formation and HA accumulation. **a)** Schematic of doxycycline-inducible expression of HAS2 in MCF10A cells. **b)** Diagram of spheroid embedding into collagen type I-based hydrogels cast into polydimethylsiloxane (PDMS) imaging chambers. Representative collagen reflectance imaging of collagen fibrillar structure (6 mg/mL). Scale bar, 50 µm. **c)** Brightfield images and quantification of collagen-embedded spheroid size after two days of culture in 2.5 mg/mL and 6 mg/mL collagen hydrogels. *indicates HAS2-dependent translucent areas. Scale bars, 200 µm. Each data point is one spheroid with n=6–17 spheroids per condition. Compact letter display is used for graphically showing statistical testing results; groups not sharing a letter are significantly different p<0.05 as calculated with a two-way ANOVA with Sidak’s multiple comparison. **d**) H&E-stained paraffin-sections of spheroids cultured in collagen hydrogels for 2 days. Scale bar =200µm and 50µm (insets). **e**) Optical coherence microscopy (OCM) images of representative collagen-embedded spheroids with and without doxycycline induction after 2 days of culture. Scale bars =200µm and 50µm (insets). Binary segmentation of the OCM slices with cellular and lumen areas displayed in green and black, respectively (left panel). 3D reconstructions from the binary images with corresponding quantification of lumen volume (right panel). **f)** Patient ultrasound image of a 0.9 cm complex cystic mass. Dotted line denotes cyst boundaries, * indicates fluid-filled luminal spaces, white arrow indicates internal septa. Scale bar =0.5 cm. Image courtesy of Dr. Victoria Mango, MD. **g)** Paraffin-sections of day 2 collagen-embedded spheroids stained with fluorescently labeled HABP. Scale bars =200µm. **h)** HA ELISA of the spheroid lysates (left) and the culture media of the suspension cultures (right). n=3 where each data point is 3 pooled spheroids or the media from 3 pooled samples, *p<0.05 from an unpaired t-test with Welch’s correction. Intraspheroidal HA concentration calculated by dividing HA content by average spheroid volume. Schematics created with BioRender.com. Data are mean ± standard deviation.

Given that the presence of translucent areas within spheroids corresponded with HA overproduction (**Fig. 1c, Extended Data Fig. 2f**), we speculated that they were cell-free lumen-like structures filled with HA. Indeed, histology confirmed the presence of cell-free regions in collagen-embedded HAS2 spheroids while parental spheroids had a uniform dense packing of cells throughout the spheroid (**Fig. 1d**). *In situ* optical coherence microscopy (OCM)^45,46^ additionally revealed that lumens were distributed throughout the full 3D volume of the HAS2-overexpressing collagen-embedded spheroids (**Fig. 1e, Extended Data Fig. 5**, axial view) and confirmed that cell-free lumens were only present when HAS2 expression was induced by doxycycline (**Extended Data Fig. 2g-h**). Quantification of lumen formation further showed that over 10% of a representative HAS2 spheroid consisted of lumens while the corresponding parental controls contained less than 0.1% (**Fig. 1e**). Interestingly, OCM images of collagen-embedded HAS2 spheroids strikingly resembled clinical ultrasound images from a patient diagnosed with a complex cystic mass similarly featuring multiple fluid-filled regions divided by internal septa (**Fig. 1f**).

To confirm that HAS2 overexpression led to increased production of HA in 3D collagen-embedded spheroids, we stained histological cross-sections with hyaluronic acid binding protein (HABP) and observed abundant HA in the spheroids and surrounding matrix of HAS2 spheroids but not parental controls (**Fig. 1g**). Because HA may have been lost during histological processing, and to quantify intraspheroidal HA accumulation vs. secretion into the surrounding media more accurately, we cultured spheroids in suspension and then performed ELISAs on both the media and spheroid lysates. First, we verified that suspension culture led to HAS2-dependent but proliferation-independent spheroid expansion and lumen formation similar to their collagen-embedded counterparts (**Extended Data Fig. 6**). ELISA next showed that HA accumulated predominantly within HAS2 spheroids as the concentration of HA contained in the lysates were approximately 800-fold higher than those measured in the surrounding media (**Fig. 1h**). In summary, HAS2 overexpression promotes intraspheroidal lumen formation, an observation that was accompanied by increased intraspheroidal HA concentration and mimicked morphological features of breast cysts in patients.

### Intraspheroidal HA accumulation elevates hydrostatic pressure within spheroids

During embryonic development, secreted HA drives lumen formation and tissue expansion by generating an osmotic gradient to create hydrostatic pressure that mechanically shapes tissues^13,17^. To test whether intraspheroidal HA accumulation correlates with elevated hydrostatic pressure^13^, we punctured the outer cell layer of spheroids using multiphoton laser ablation^47,48^(**Fig. 2a**) and monitored for outward fluid flow, indicative of increased pressure^49^. For these experiments, we fluorescently labeled spheroids and cell culture media, ablated microtrack^50^ channels in the collagen immediately adjacent to each spheroid, and ablated the spheroid’s outer cell layer to allow outward fluid flow from the spheroid into the channels to be visualized (**Fig. 2a**). FITC in the media and fluid contained in the HAS2 spheroids (**Extended Data Fig. 7a**) provided contrast with the non-fluorescent, liquid-free cavitation bubble that formed in the channels during laser ablation. We quantified the magnitude and direction of fluid flow by tracking the displacement of the cavitation bubble and the motion of fluorescently labeled cellular debris (**Fig. 2b-f**) through the pores ablated in the outer surface of each spheroid. Immediately after ablation, medium flowed outward from HAS2 spheroids and displaced the cavitation bubble unidirectionally, whereas parental spheroids showed little outward flow, resulting in slower, more uniform bubble displacement and medium flow into the spheroid (**Fig. 2c, green triangle**). Over the short 120 s duration of the videos, HAS2 spheroids showed a trend toward greater bubble displacement, but the difference was not statistically significant. Over a longer timescale, however, flow could be measured more accurately: phase contrast images acquired after 1 hour showed significantly greater displacement of cell debris into the microtracks for HAS2 versus parental spheroids (**Fig. 2d**), demonstrating sustained outward flow under these conditions. To estimate the pressure differential across the spheroid surface^47,48^ we quantified the velocity of cell debris (**Fig. 2e-f**) and used the Hagen–Poiseuille equation to model fluid flow through the ablated pore as pipe flow (**Fig. 2g, Extended Data Figure 7b,** accounting for the radius of each independent pore in the outermost layer of the spheroid) to estimate a pressure difference of 8 Pa ± 5.5Pa (SD) in the HAS2 spheroids (**Fig. 2h**)^51^. Collectively, these data suggest that intraspheroidal accumulation of HA promotes cyst-like lumen formation in HAS2 spheroids via the generation of hydrostatic pressure^48,51^, a finding that is consistent with prior work on hydraulic lumen inflation in Madin-Darby canine kidney type II (MDCK-II) cysts^48^ and pancreatic organoids^49^.

**Figure 2:**
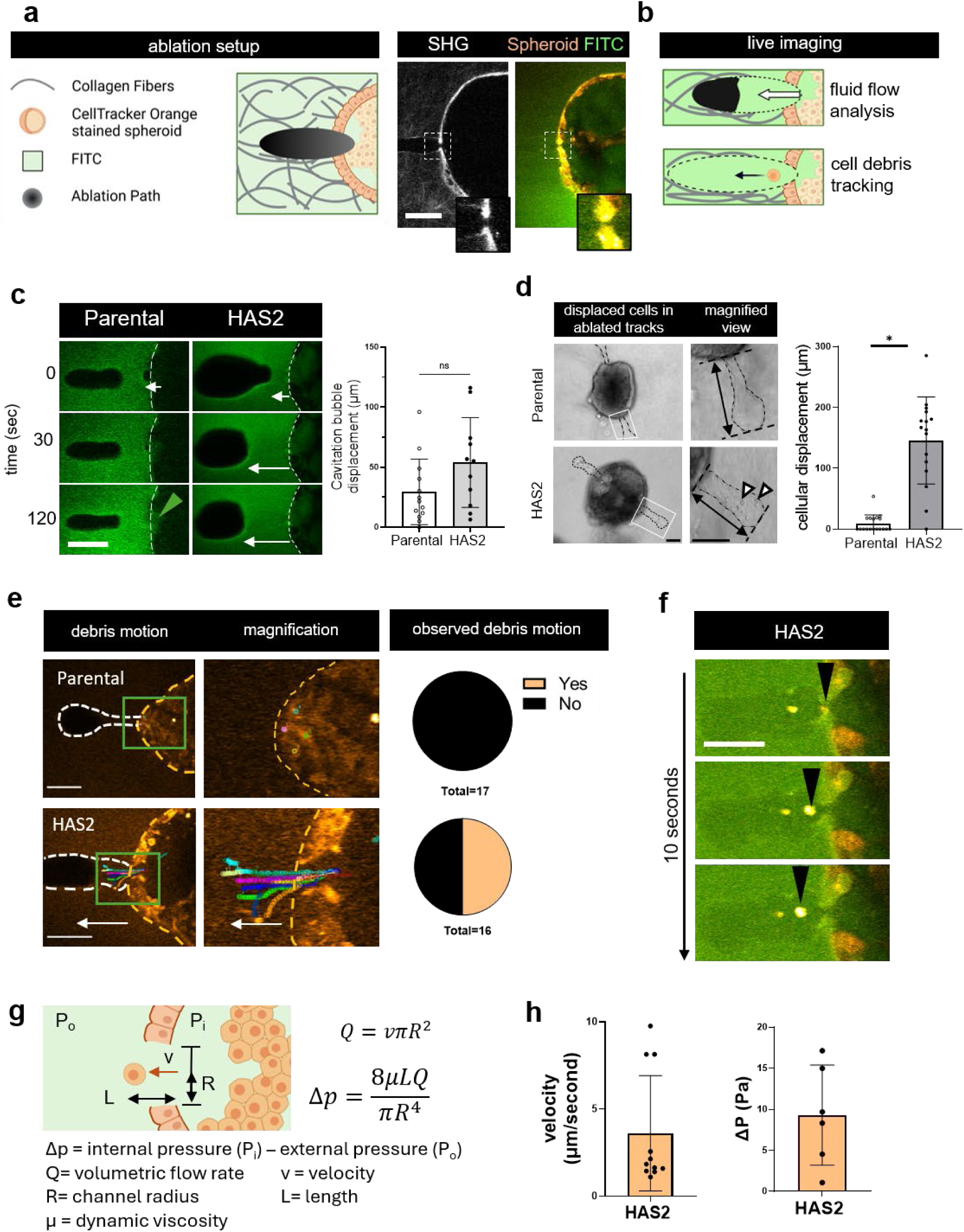
Laser ablation reveals HAS2 overexpression generates intraspheroidal hydrostatic pressure. **a)** Schematic depicting the multiphoton laser ablation experiments. The microtrack ablation process generated a cavitation bubble which filled the ablated space until it was displaced by fluid. Representative images show the collagen SHG signal (grey) and the CellTracker-stained spheroid (orange) with FITC labeled media (green). Insets display the ablated gap in the outer densified collagen and outer cell layer of the spheroid. Scale bar =100µm. **b)** Schematic of the analytical pipeline used to estimate hydrostatic pressure from fluid flow, quantified by tracking cavitation bubble displacement and cell debris transport following laser ablation. **c)** Cavitation bubble (black region) displacement by FITC labeled cell culture media (green). White arrows indicate fluid flow over time, orange dotted line demarcates the spheroid boundary, the green triangle indicates media diffusing into the parental spheroid. Bubble displacement from the cell-ECM interface is quantified over 120 seconds. Each data point represents one ablation track. n=13 parental, n=11 HAS2. No statistical difference of *p<0.05 as a determined by an unpaired t test with Welch’s correction. **d)** Phase contrast images of spheroids taken after ablation with the ablated tracks annotated in black and displaced cellular debris highlighted with white triangles. Scale bars =100 µm. The maximum distance of displaced debris in each track is increased in HAS2 spheroids suggesting outward flow. Each data point represents one ablation track, n=16-18 tracks from 8-9 spheroids *p<0.05. Statistical significance determined by an unpaired t-test with Welch’s correction. **e)** Colored tracks, generated with the MTrackJ plugin in Fiji, show the motion of individual pieces of cell debris over time. White arrows indicate the direction of cell motion. Each color track is an independent particle; scale bar =100µm. **f)** Representative image of a labeled cell passing through the ablated opening. Scale bar =50µm. **g)** Schematic of the parameters measured from the ablation videos and used to calculate particle velocity and, with the Hagen-Poiseuille equation, to estimate the pressure difference between the spheroid lumen and the surrounding environment. **h)** Quantification of particle velocity (1-2 particles quantified per track; each point is one particle). Quantification of the estimated pressure difference between the inside and outside of the spheroid. Each data point represents one track. Schematics created with Biorender.com.

### Expanding HAS spheroids generate mechanical compression at the cell-ECM interface

Benign lesions are often surrounded by collagen fibers that are aligned parallel to the tumor boundary (often referred to as tumor-associated collagen signature 2, TACS-2^28^), and these changes have been attributed to compressive forces caused by lesion growth^52^. To determine if HA overproduction also causes elevated mechanical stress at the interface between the spheroid and surrounding collagen, we co-embedded fluorescent deformable polyacrylamide beads into the collagen gels as *in situ* pressure sensors^53–57^ (**Fig. 3a**). Beads located at the spheroid-ECM interface exhibited greater anisotropic deformation than those localized in the bulk collagen (**Fig. 3a-b**), suggesting that HAS2-overexpressing spheroids created pressure at the surrounding collagen interface.

**Figure 3:**
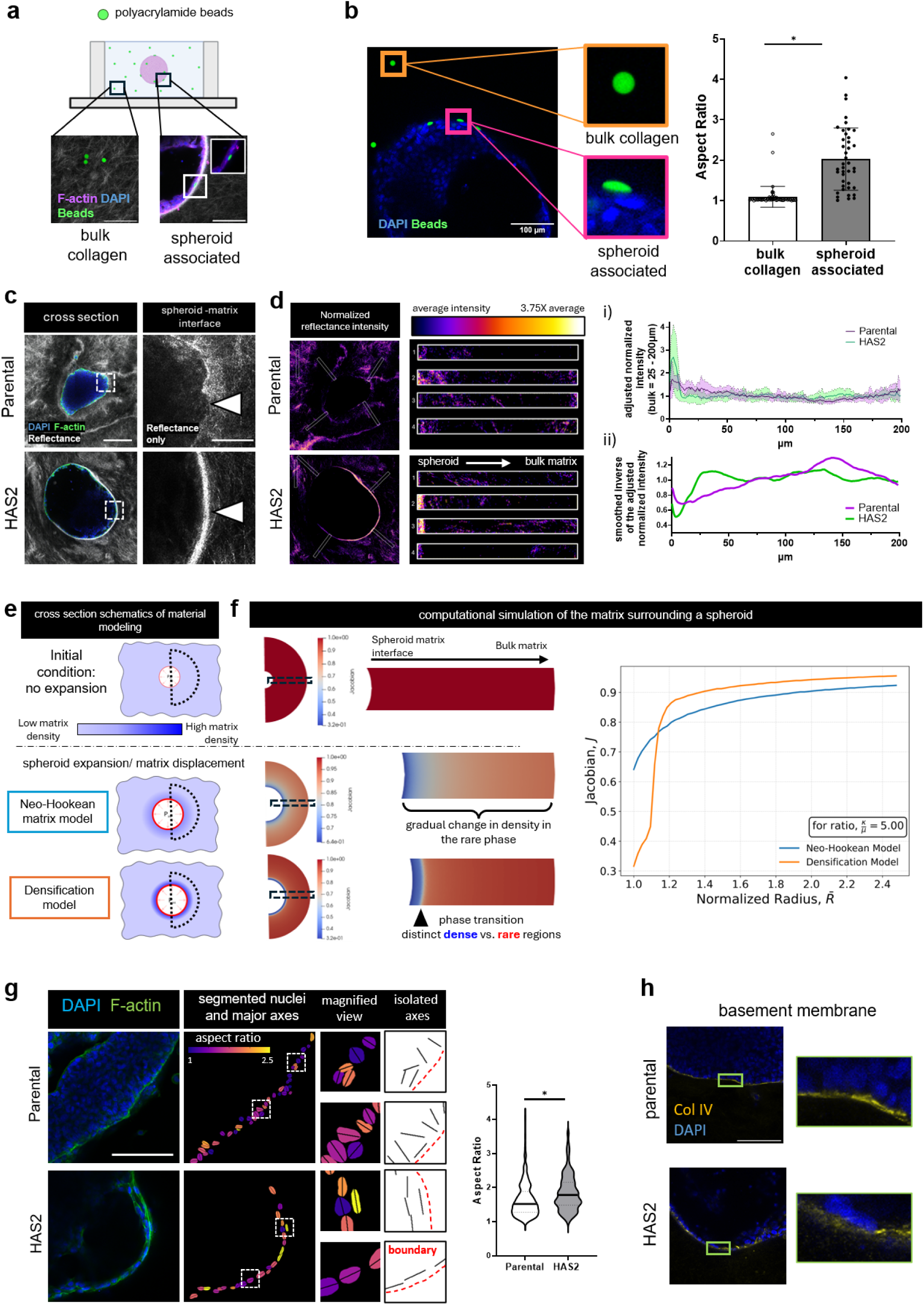
Spheroid expansion creates altered biomechanical forces at the spheroid-matrix interface. **a)** Schematic of polyacrylamide beads incorporated into the spheroids and the surrounding ECM as *in situ* pressure sensors and corresponding representative confocal images of beads in bulk collagen or associated with spheroids. Scale bar = 100 µm. **b)** Confocal images of representative spherical beads in the bulk collagen (upper orange inset) and elongated beads at the spheroid-ECM interface (lower pink inset) and corresponding quantification of bead aspect ratio. Scale bar = 100 µm. Each data point represents one bead; n=61 outside, n=41 spheroid associated; * = p<0.05 as determined by an unpaired t-test with Welch’s correction. **c)** Confocal and collagen reflectance images of day 3 spheroids revealing differences in collagen densification between parental and HAS2 spheroids (white arrows). Scale bar = 200µm and 50µm (magnified insets). **d**) Corresponding collagen reflectance intensity images (normalized to bulk intensity) with 4 radial inserts (white boxes) displaying collagen intensity as a function of distance from the spheroid-matrix interface (plot i). Plot ii) shows the inverse of the average collagen intensity smoothed with a 20-point moving average filter for comparison to computational results shown in f). **e)** Overview schematics of finite element simulation and results of a pressurized spheroidal cavity expanding into a foam-like compressible material representing the ECM. The red to blue gradient corresponds to low and high matrix density. Black semi-circles denote the matrix regions analyzed in (f). **f)** Finite element simulations of a compressible foam-like hyperelastic material using FeniCS finite element open-source library. A spherical solid of outer radius R=5, and inner cavity with radius r=1 is pressurized internally. The quantity plotted is the Jacobian of the deformation gradient which indicates the volume ratio at every point in the body. J(x)=V_final_/V_initial_ indicating the ratio of final to initial volume pointwise in the continuum. An increase of the Jacobian corresponds to a decrease in concentration (per deformed volume of the body) of the solid phase of the material. **g)** Confocal images of nuclear morphology at the spheroid-ECM interface. Segmented nuclei are color-coded by aspect ratio. The major axis of each nucleus was calculated, overlaid in black, and displayed independently in the rightmost insets with the spheroid boundary drawn in red. Violin plots of nuclear aspect ratios in parental and in HAS2-overexpressing spheroids. Each data point is one nucleus from multiple depth planes in 3 spheroid from each condition *= p<0.05 as determined by a Mann-Whitney test, n=270 parental, n=168 HAS2. **h)** Collagen IV (yellow) staining of spheroid boundary illustrates the presence of a basement membrane in both conditions. Scale bar =100µm. Schematic in e) created with BioRender.com.

Prior work has shown tumor cell proliferation within a spheroid induces compression and densification of the surrounding matrix via collagen fiber bending^52^. In our case, spheroid expansion arises from HA-driven hydrostatic pressure rather than by cell proliferation alone. Collagen densification (**Fig 3c**, white arrows) at the matrix interface of control spheroids was evident, but markedly less pronounced than the dense band surrounding HAS2 or HAS3 spheroids (**Extended Data Fig. 3f-g**). An analysis of radial matrix profiles from the spheroid surface extending 200µm out into the bulk collagen revealed a sharp transition from dense collagen at the interface of HAS2 spheroids to the looser fiber structure of the bulk matrix (**Fig. 3d**). This localized densification response is reminiscent of the behavior of compressible foams where phase transition signatures emerge to reveal a sharp boundary between the highly compressed proximal region (dense phase) and the distal low-density region (rare phase). This phase transition is observed in fibrin^58^ and collagen^37,59^ networks undergoing linear uniaxial compression. The emergence of distinct phases contrasts with the response expected from a compressible biopolymer network without phase transitions where a smooth decrease in collagen density is expected. We used finite element analysis to model our unique geometry by simulating pressurized cavity expansion in a system representative of the ECM (**Fig. 3e, Supplementary Note 1**). This allowed us to test whether our experimental findings were consistent with constitutive models that do or do not support phase transitions and abrupt volumetric densification. Specifically, we sought to compare the response of the Helmholtz free energies between a Neo-Hookean (polyconvex) model that prevents phase transition and a densification (non-convex/multi-well) model that permits phase transitions for the volumetric response^38^. Finite element simulation results demonstrate a loading-dependent (**Extended Data Fig. 8**) response of material densification suggesting that the extent of spheroid expansion dictates collagen densification. Radial regions of interest starting from the simulated spheroid-matrix interface show a continuous density gradient in the Neo-Hookean model and the expected presence of a discrete density transition at a finite distance from the spheroid-matrix interface in the densification model (**Fig. 3f**). Correspondingly, the Jacobian (ratio of current to initial volume pointwise) plot reveals the phase transition in the densification model as identified by the characteristic steep slope capturing densification (**Fig. 3f**). These computational results further support the hypothesis that spheroid expansion driven by HA-dependent hydrostatic pressure within the spheroid compress the surrounding collagen to promote a local phase transition in the ECM creating a band of densified collagen at the spheroid-matrix interface.

While the formation of a physical barrier due to collagen phase transition could limit invasion independently, the morphology of the cells at the spheroid-ECM interface also differed between conditions. In HAS2 conditions, cells immediately interacting with the surrounding collagen formed a single monolayer of cells with elongated nuclear morphologies and circumferential orientation, potentially indicating a more myoepithelial cell phenotype (**Fig. 3g**) as opposed to the dense, more spherical and columnar orientation of nuclei found in the parental spheroids suggestive of a more luminal epithelial cell phenotype (**Fig. 3g**). Collagen compaction and circumferential cell orientation findings were also present in lower density collagen hydrogels (**Extended Data Fig. 9**) further supporting that expansive forces caused by intraspheroidal HA accumulation may create hoop stress that results in elongation and tension-driven alignment of cells in immediate contact with collagen^60^, but not excluding the possibility that the observed cellular orientation was promoted by spheroid rotation^61,62^. To further interrogate the spheroid-matrix interface, we stained for the basement membrane protein collagen IV ^63^ (**Fig. 3h**). While both spheroids assembled collagen IV, parental spheroids featured more organized collagen IV staining. This suggests that HA overproduction may alter basement membrane formation implying differences in invasive potential^63^. Collectively, our findings indicate that HA-driven lumen formation and subsequent spheroid expansion generate compressive forces on the surrounding matrix that contribute to collagen densification at the spheroid interface and regulate cell alignment in the outer monolayer, which, together with localized matrix alignment at the spheroid interface^61,62,64^, may synergistically inhibit invasion.

### Computational modeling suggests osmotically driven paracellular fluid flow as a mechanism for lumen formation and localization

To better understand the biophysical mechanisms by which HA may drive lumen formation and intraspheroidal hydrostatic pressure, we developed an *in silico* model^65^ in which we could selectively adjust cell-cell adhesion strength and paracellular fluid flow. We employed an experimentally validated cell-based vertex model^66,67^ in which cells share interfaces within a two-dimensional representation of a spheroid cross-section^65^ (**Supplementary Note 2**). Cells are modeled as deformable polygons that are energetically regulated by bulk elasticity (energetic cost for change in area) and contractility of the actomyosin cell cortex (energetic cost for change in perimeter) (**Fig. 4a**). Cell-cell interfaces are separated by a gap to allow for a paracellular transport-inspired^13,48^ flow network modeled as fluid flow through a pipe. The distance between cells (pipe width) is modeled as springs with tunable spring constants allowing HAS-dependent changes in MCF10A cell-cell adhesion receptor expression^68^ and the resulting adhesion strength to be incorporated into the model. Taken together, paracellular fluid transport into the spheroid is modeled as pipe flow regulated by 1) pipe width (which is inversely related to the strength of cell-cell adhesions) and 2) differences in osmotic pressure (which drives fluid flow to regions of high HA concentration (**Fig. 1h**) to generate quantifiable hydrostatic pressure (**Fig. 2h**)) (**Fig. 4a**). We call this novel version of tissue an extended vertex model with a paracellular flow network.

**Figure 4:**
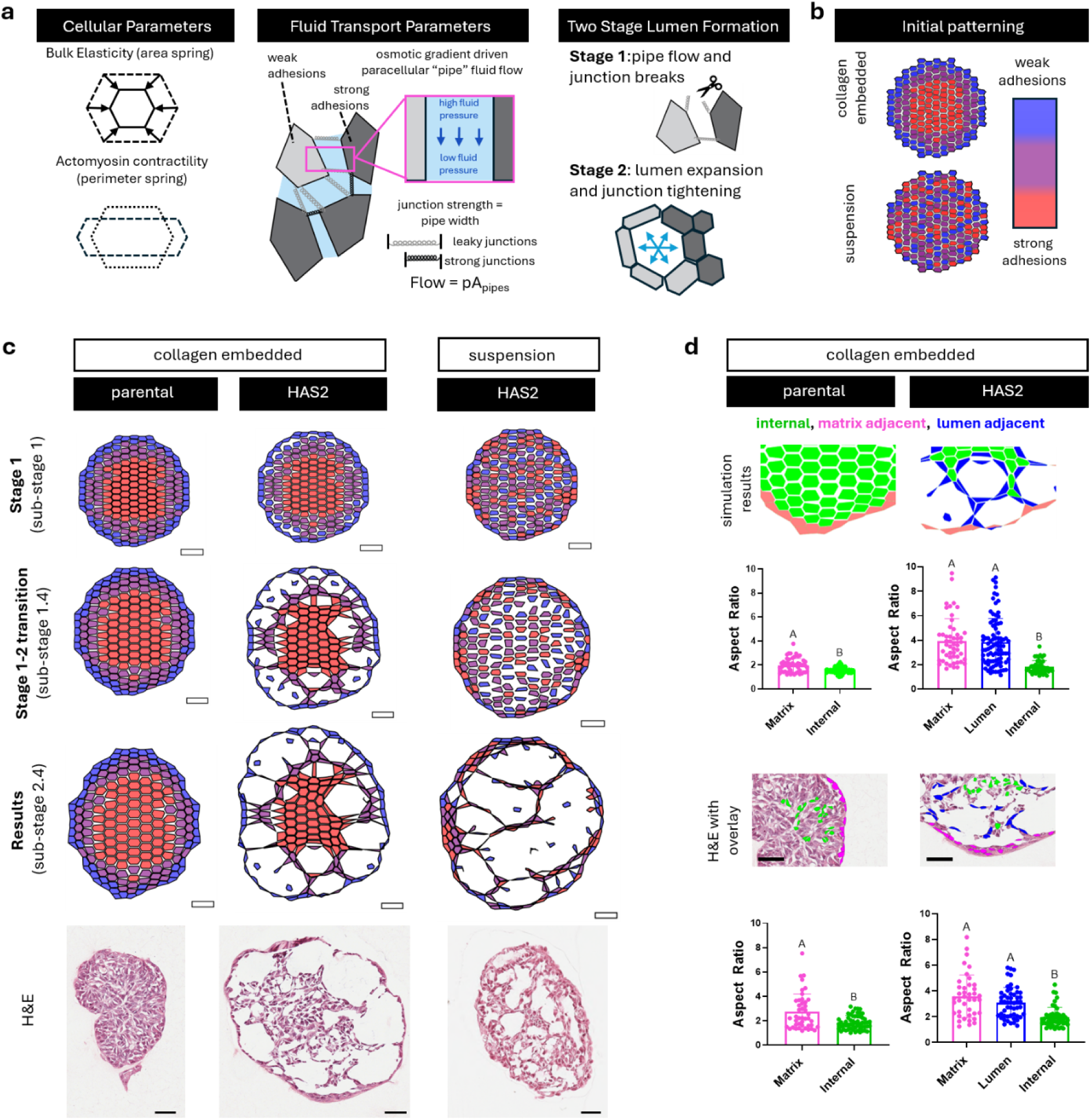
Computational models demonstrate that lumen formation is a function of osmotic pressure and intraspheroidal lumen location is determined by the spatial distribution of cell adhesion strengths. **a)** Schematic depicting the cellular and fluid transport parameters of the *in silico* model of 2D spheroid cross-sections. The two-stage model of lumen formation highlights the actin remodeling, pipe flow, and breaking of cell-cell adhesions that occur in stage 1 and the osmotic pressure-driven lumen formation that occurs in stage 2 when the remaining cell-cell adhesions tighten. **b)** Initial configuration of collagen embedded parental and HAS2 spheroid cross-sections where cells with strong cell-cell adhesions (red) cluster in the center and cells with weak cell-cell adhesions (blue) are located at the periphery of the spheroid as predicted by the differential adhesion hypothesis (Supplementary Note 2). In the absence of confinement-induced spatial patterning, we assume a uniform distribution of cell adhesion strengths in the cross section of a suspension culture spheroid. **c)** The simulation results and H&E images of experimental samples after 2 days display similar bulk structural morphologies with lumens only present in HAS2 conditions in the model where there is an osmotic pressure gradient and reduced cell-cell adhesion strengths. HAS2 suspension culture is simulated with non-sorted cells in the initial condition. Scale bars are 4 A.U. = 50µm in the simulations and scale bars = 50μm in the H&E images. **d)** Insets of regions at the cell-matrix interface in the simulations and experimental samples annotated with cell traces (green=internal cells, pink = matrix adjacent cells, blue = lumen adjacent cells) used to quantify morphological differences in cells as function of their location in the spheroid. Each data point represents 1 cell with the number of cells listed as n = matrix, internal, lumen. Parental simulation n= 51,141. HAS2 simulation n= 50, 47, 93 (6 lumen outliers excluded). Parental experimental n=44, 57. HAS2 experimental n=40, 48, 52. Statistical significance as determined by p<0.05 via an unpaired t test with Welch’s correction for the parental conditions and a Brown-Forsythe and Welch ANOVA test with multiple comparisons for the HAS conditions. Statistics displayed with compact letter notation. H&E scale bars =50μm.

To simulate lumenogenesis in our model, we implemented a multiple stage approach in which lumen formation is initially driven by actin remodeling (stage 1) followed by osmotic pressure-driven lumen expansion (stage 2) as motivated by prior experimental work^51^. Stage 1 in our model was characterized by simulated actin-driven cell tension, the breaking of highly strained cell-cell adhesions (spring junctions), and simulated fluid flow through the pipe network, set as function of water movement in response to HA accumulation-dependent osmotic gradients. During stage 2, the remaining cell-cell adhesions (springs) tighten^51^, and sites of broken cell-cell junctions may become lumen initiation sites that expand through osmotic gradient-driven simulated fluid influx (**Fig. 4a**). Because collagen-embedded, HA-overproducing spheroids formed lumens near the spheroid periphery, whereas lumens in suspension-culture spheroids were predominantly centralized (**Extended Data Fig. 6c**), we next leveraged our model to determine how collagen confinement regulates the spatial localization of lumen initiation. Consistent with our experimental findings (**Supplementary Note 2, Fig S2**), we assumed in these simulations that HA-overproducing spheroids in suspension culture contained a homogeneous mix of cells with weak and strong cell-cell adhesions, whereas their counterparts embedded in collagen exhibited spatial patterning of cell populations according to adhesion strength, a process known to be triggered by mechanical confinement^69^ (**Fig. 4b**). Details of the modeling equations, input parameters, and biological justification can be found in Supplementary Note 2.

Strikingly, collagen-embedded simulated parental spheroids had no lumens, while their simulated HAS2-overexpressing counterparts exhibited peripherally localized lumens (**Fig. 4c**), which grew as a function of simulated time (**Extended Data Fig. 10a**). The lack of lumens in the parental condition was controlled by either the absence of osmotic gradient-driven fluid influx or sufficiently strong cell-cell adhesion that resisted rupture, thereby preventing the formation of robust pipe flow (**Extended Data Fig. 10b**). These results support a fluid-flow-driven mechanism of lumen formation and explain the different localization of lumens in collagen-embedded vs. suspension culture spheroids (**Fig. 4c**). The presence of floating cells in the lumens of simulated spheroids is also consistent with our experimental observations (**Fig. 2**) and corresponds to a HAS-induced decrease in E-cadherin expression^68^ and thus, reduced cell-cell adhesion potential (**Supplemental Note 2, Fig S2**). Finally, simulated cells adjacent to either the spheroid-matrix interface or a lumen exhibited an elongated morphology similar to our *in vitro* results further validating the capability of our extended vertex model to reveal new mechanistic insights into cyst-like lumen formation (**Fig. 4d**). The physical deformation of cells further suggests phenotypic heterogeneity within the spheroid induced by spatial variation in mechanical forces and recapitulates our experimental findings, which identified nuclear elongation in the outer layer of HAS2 spheroids suggestive of elevated surface tension adjacent to the pressurized lumens (**Fig. 3g**).

In summary, our extended vertex model incorporating a paracellular flow network uniquely enables the study of lumenogenesis sites within spheroid cross-sections, without requiring these sites to be predetermined, as was necessary in previous models^51^. Using the model, we found that lumen position depended only on the final arrangement of cells at the onset of modeled junction rupture at the end of stage 1 and not on the kinetics of adhesion-based cell sorting (**Extended Data Fig. 10**). Moreover, our model suggests that HAS-overexpressing spheroids exhibit both time-dependent hydraulic lumen growth driven by the accumulation of osmotically active HA^70,71^ and spatially regulated lumen nucleation^72^ determined by matrix confinement and adhesion-mediated cell patterning.

### Adipose stromal cells (ASCs) suppress lumen formation and promote spheroid invasion through ECM remodeling

Our results suggest that cyst-like intraspheroidal HA accumulation inhibits invasion by promoting lumenogenesis-induced compression and densification of the surrounding ECM (**Figs. 1-4**). Indeed, elevated ECM density and solid stress have previously been shown to decrease tumor invasion and growth^73,74^. In that work, however, ECM compression arose from highly contractile cancer-associated fibroblasts (CAFs) rather than from a hydrostatic pressure-induced ECM phase transition. Moreover, CAFs promote rather than inhibit cancer invasion in many other contexts, and ECM remodeling is critical to this process^75–77^. In the context of breast cancer, CAFs can emerge from ASC differentiation into myofibroblastic CAF-like phenotypes and mediate tumor cell invasion via ECM remodeling, the formation of direct heterotypic cell-cell adhesions, and soluble factor signaling^78,79^. To better understand how stromal cells affect cyst-like lumen formation, expansion and invasion, we incorporated ASCs into the collagen surrounding HAS2-overexpressing MCF10A spheroids at a 6:1 cell ratio. This ratio was chosen to mimic an elevated population of stromal cells in the surrounding stroma when heterotypic cell-cell adhesions are rare and only localized at the spheroid periphery (**Fig. 5a**). Consistent with previous findings, ASCs in the surrounding collagen did not promote outward invasion from the spheroids^32^, and failed to prevent lumen formation and spheroid-adjacent ECM compression, suggesting that soluble factor signaling between ASCs and HAS2-MCF10A cells was insufficient to promote the shift to an invasive phenotype (**Fig. 5a**). To better mimic *in vivo* scenarios in which ASCs directly interact with tumor cells, co-culture spheroids were formed by mixing HAS2-overexpressing MCF10A cells and ASCs at a 1:1 ratio (**Fig. 5b**). Embedding co-culture spheroids into collagen revealed that ASCs in direct physical contact with MCF10A cells markedly increased invasion as compared to incorporating cells into the bulk collagen regardless of HAS2 expression (**Fig. 5c-e**). Both types of co-culture spheroids featured both radial collective and single-cell invasions led by alpha smooth muscle actin (αSMA)-positive ASCs (**Fig. 5c**). In contrast, neither mode of cell migration was observed in monocultures or co-cultures with ASCs in bulk collagen (**Fig. 5e**). Peripherally located luminal structures were noticeably lacking in the HAS2-co-culture spheroids, a finding perhaps explained by the fact that the ASCs^80^ interfere with the formation of cell-cell adhesions between HAS2-MCF10A cells and therefore prevent HA accumulation and lumen formation. To quantify ASC-mediated differences in spheroid shape, we used an average radial length crossing quantification,^81^ adopted from mammography analysis^82^, as a metric of border irregularity and collective invasion. This approach revealed that co-culture spheroids had significantly more irregular borders (**Fig. 5e**), a characteristic associated with lesion malignancy and worse prognosis in the clinic^83^. Analysis of the ECM surrounding the co-culture spheroids revealed the absence of densified collagen that we previously observed for monoculture spheroids, suggesting ASCs prevent ECM compression at the spheroid borders (**Fig. 5d**). Given these observations, we hypothesized that ASC-mediated ECM remodeling was critical to overcoming the invasion-inhibiting effect of collagen compression that was mediated by intraspheroidal HA accumulation. To test this hypothesis, we broadly inhibited matrix metalloproteinases (MMPs) using batimastat in HAS2 co-culture spheroids. Inhibiting MMPs prevented ASC-mediated invasion, rescued the phenotype of collagen densification at the spheroid borders, and prevented spheroid expansion and the formation of observable lumen structures (**Fig. 5f**), suggesting that ASCs promote invasion by preventing HA accumulation-dependent spheroid expansion and collagen compression. To rule out potential confounding effects of batimastat treatment on HAS2-overexpressing MCF10A cells, a subset of monoculture spheroids was also treated. While batimastat reduced the expansion of monoculture spheroids, it did not affect lumen formation (**Extended Data Fig. 11a**). Because ASCs were α-SMA positive, indicative of a contractile phenotype (**Fig. 5c**, green), we also inhibited contractility through blebbistatin treatment. This treatment did not prevent the invasion of co-culture spheroids and correspondingly no collagen densification was present at the spheroid matrix interface (**Extended Data Fig. 11b-c**), indicating the contractile phenotype of the stromal cells was not driving the physical matrix remodeling required for invasion. In summary, invasion of HAS2 overexpressing MCF10A cells from spheroids was only observed in co-culture conditions where MMP-expressing ASCs made direct contact with epithelial cells. These heterotypic interactions likely disrupted epithelial cell-cell adhesions and prevented the accumulation of intraspheroidal HA and hydrostatic pressure to drive spheroid expansion. Disruption of lumen formation combined with the matrix remodeling capabilities of the ASCs prevented localized collagen densification via phase transition and allowed HAS overexpressing cells to invade into the surrounding matrix along with the ASCs (**Fig. 5g**), a finding which matches single cell invasion data^8^.

**Figure 5:**
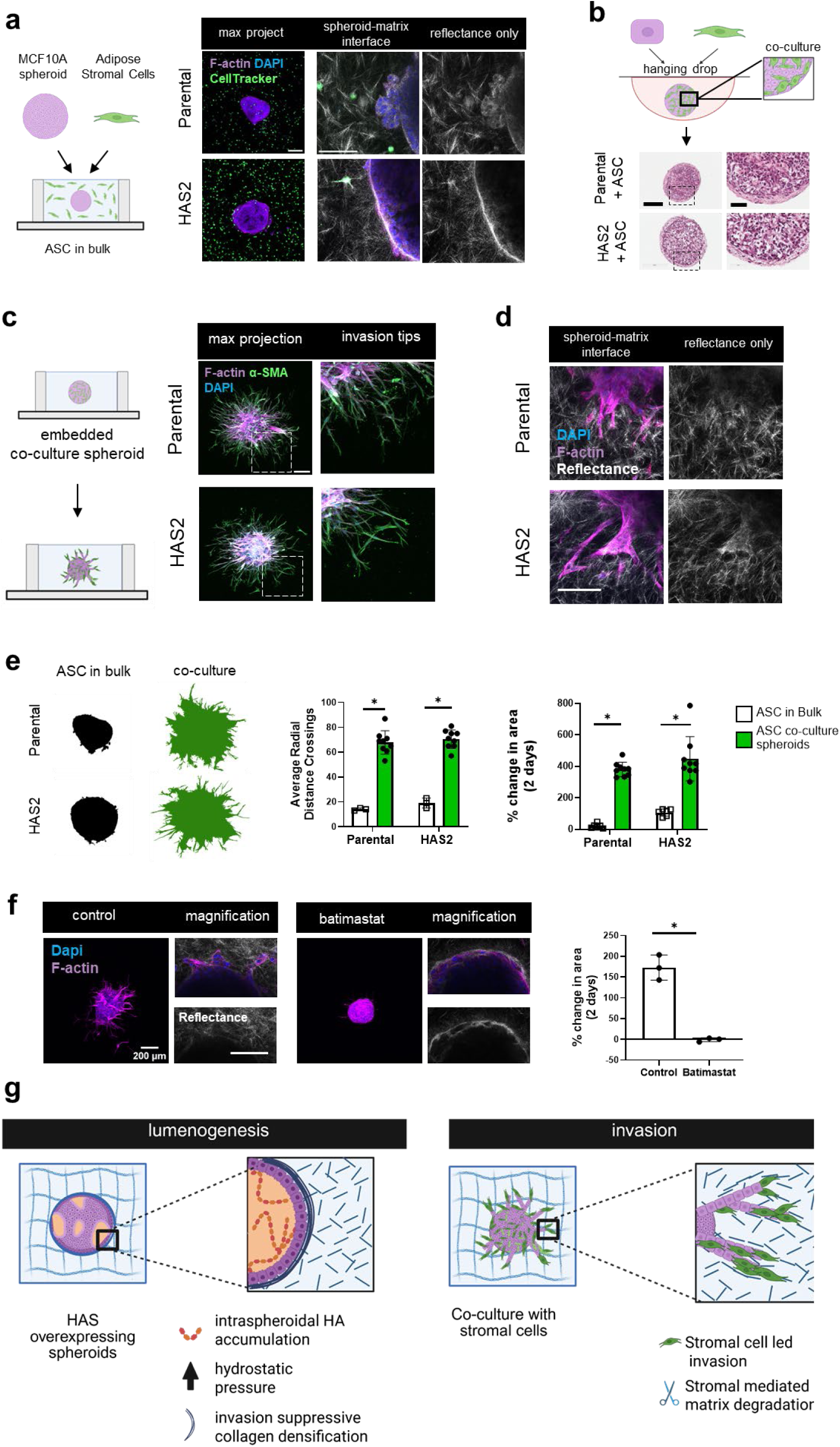
ASCs enable invasion of HA-overproducing MCF10A spheroids by interfering with collagen compression and densification. **a)** Schematic of CellTracker green labeled ASCs in the bulk collagen ECM surrounding an MCF10A spheroid and representative confocal micrographs of day 2 cultures of parental or HAS2 MCF10A spheroids. CellTracker labeled ASCs in green with all cells stained for DAPI (blue) and F-actin (magenta) and collagen reflectance in grry. Scale bars =200 µm. **b)** Schematic of co-culture spheroid formation where ASCs and MCF10A cells are mixed in a 1:1 ratio in hanging drops. H&E images demonstrate cell-cell contacts and mixing of cell types. Scale bar =200µm (upper image) and 50µm (lower zoomed in images). **c)** Schematic of co-culture spheroid embedded into collagen hydrogels. Maximum confocal projections after a 2-day culture period in collagen hydrogels show radial collective and single-cell invasion. Cells stained with DAPI (blue), F-actin (magenta), and αSMA (green). Magnified images of invasion tips show α-SMA positive ASCs lead invasion strands. Scale bar =200µm**. d)** Confocal slices of F-actin (magenta) stained invasion boundaries and collagen reflectance imaging (grey) show an absence of a densified collagen surrounding the spheroid. Scale bar=100µm. **e)** Outlines of collective spheroid invasion were generated as regions of interest from maximum projections of confocal images and used for analysis of surface irregularities by analyzing average radial distance crossings and invasion area. Each data point is one spheroid with n=3 for bulk ASC co-cultures and n=9 for the direct ASC co-cultures. **f)** Maximum projections of ASC-HAS2-MCF10A co-cultures with and without the MMP inhibitor batimastat stained with phalloidin (magenta) and DAPI (blue). Slices display collagen reflectance in grey, scale bar, 100µm. Quantification of percent change in area of spheroid invasion, each data point is one spheroid. All statistics are unpaired t-tests with Welch’s correction with *p<0.05. Schematics created with BioRender.com. **g)** Schematic of suggested model of how HA overproduction and intraspheroidal accumulation inhibits invasion by driving cyst-like spheroid expansion and collagen compression. Direct cell-cell contact with ASCs can overcome both effects to enable invasion into the surrounding ECM. Schematic created with BioRender.com.

### Outlook

Although HA is recognized as a mechanically active biomaterial in physiological, therapeutic, and tissue engineering contexts^3^, its role in breast cancer has been viewed largely through biochemical and metabolic activity^8,10,84^. Here, we built physiologically relevant 3D models which can recapitulate tissue-level HA accumulation permitting previously inaccessible insights into how HA physically regulates the breast. Our results suggest that epithelial confinement of secreted HA forms osmotically active biomaterial pockets within epithelial lesions that initiate hydrostatically driven lumen formation to mirror key features of embryonic lumenogenesis^13,16,17,85^ and HA-driven cystic cancers ^18,19,86^, producing spheroid architectures that closely resemble benign multi-luminal complex mammary cysts^87^. Critically, expanded spheroids mechanically compress the surrounding collagen to create a dense invasion-suppressing matrix by driving collagen phase transition^37,52^ at the spheroid-matrix interface. Collectively, our findings suggest that HA regulates breast tissue fate through currently underappreciated biophysical mechanisms that act alongside, and potentially counterbalance, its known biochemical signaling and metabolic effects.

Although collagen architecture is known to regulate solid tumor progression^64^, it is unclear whether HA swelling, combined with the matrix mechanics and structural changes characteristic of fibrocystic breast disease, helps determine if a breast lesion progresses to malignancy or remains benign. Fibrillar collagen matrices exhibit complex mechanical behavior, including strain stiffening, compression softening, and biphasic material properties^15,37,88^. Our FEA results suggest localized deformation and phase transitions at the compression interface, like those observed in cellular materials^38^. Consistent with observations in other fibrillar materials, this phase transition likely arises from fiber buckling, densification, and compression-induced realignment^37,52,58,59^. Previous studies have shown that incorporating uncrosslinked HA into collagen hydrogels does not significantly alter bulk material properties. However, HA suppresses fiber densification and reduces compression softening^15,88^. This effect has been attributed to HA-induced matrix swelling, which places collagen fibers under pre-tension. These findings highlight an important area for future experimental and computational investigation, particularly when comparing monocultures containing HA within lumens to co-cultures, where HA is hypothesized to be released into the surrounding matrix.

Our *in silico* model provides a mechanistic explanation for confinement-dependent lumen distribution in spheroids, linking it to the interplay between cell-cell adhesion strength, osmotic pressure, and paracellular fluid flow. While this model was applied specifically to study breast cyst development, complementary analyses can model embryonic lumenogenesis and tissue morphogenesis^65^. Our spheroid cross-section model could be strengthened by including defined apical-basal polarity of each modeled cell^89,90^, hydraulically gated oscillations^70,89^, or lumen stability^90^ parameters. Cumulatively, models of the matrix and spheroid both suggest cells on the periphery of the spheroid experience matrix confinement-induced compression, lateral forces that elongate and stretch the cells, and induced curvature^91,92^. These forces likely activate mechanobiological modulation of cell phenotype^93^ and should be the focus of future studies.

The breast tumor microenvironment comprises more than the epithelial lesion and its surrounding matrix. Diverse stromal cells, including adipose stem cells, fibroblasts, immune cells, and adipocytes critically shape malignant transformation and tumor behavior^94,95^. Invasion was observed only when matrix remodeling stromal cells were in direct contact with the spheroid. Thus, we hypothesize that extensive MMP-mediated stromal remodeling is necessary for HAS-overexpressing benign mammary cysts to develop into invasive disease. HA-mediated signaling promotes stromal differentiation into a pro-remodeling myofibroblast-like phenotype while simultaneously increasing HA production, creating positive feedback loops within the stroma ^96^. Although immune cells are not included in the current model, incorporating macrophage or T-cell subsets could reveal whether HA-driven and stroma-driven remodeling differentially regulate immune infiltration^97,98^. Such models would also enable investigation of how an HA-rich glycocalyx modulates heterotypic interactions among immune, stromal, and tumor cells to influence disease progression ^99,100^. Taken together, our *in vitro* and *in silico* platforms enable future investigations into the role of HA in breast cyst formation and the transition to malignant invasion in microenvironmental contexts of increased stromal complexity to address unresolved questions in breast health.

## Methods

### Cell culture

Tetracycline-inducible HAS2 and HAS3 MCF10A cell lines were generated via lentiviral transduction as previously described ^41^. MCF10A cells were cultured in DMEM/F12 (Gibco 12800082/21700075) supplemented with 5% Horse Serum (Gibco 16050122), 1% penicillin/streptomycin (Gibco 15140122), 10 µg mL^−1^ Insulin (Sigma 91077C), 0.5 µg mL^−1^ Hydrocortisone (Sigma-Aldrich H4001), 100 ng mL^−1^ Cholera toxin (Krackler Scientific 45-C8052), and 20 ng mL^−1^ human epidermal growth factor (EGF, Gibco PHG0313). Supplemented DMEM/F12 referred to here as enriches DMEM/F12. Human Adipose-Derived Stem Cells (Lonza, PT-5006) (referred to as ASCs in this manuscript) were cultured in Lonza Human Adipose Derived Stem Cell Growth BulletKit Medium (Lonza PT-4505) and subcultured as recommended by Lonza. Co-culture studies were carried out in a 1:1 mix of MCF10A and Lonza media. Experiments were conducted in cell type matched media supplemented with 1 µg mL^−1^ doxycycline (Santa Cruz Biotechnology, sc-204734A) to induce HAS overexpression. Doxycycline was introduced after experiments were seeded and spheroids were formed. Co-culture spheroids were cultured at a 1:1 ratio of MCF10As:ASCs. Experiments with stromal cells in the bulk collagen were cultured in a ratio of 1:6 MCF10A to ASCs. All cells were cultured at 37°C and 5% CO^2^.

### 2D Cell culture

For 2D imaging experiments, cells were cultured on fibronectin (Gibco 33016015, 30 µg mL^−1^) or type 1 rat tail collagen (Corning 354249, 30 µg mL^−1^) coated 18mm glass coverslips. Cells were seeded overnight to adhere and induced with doxycycline the following day. Transwell migration assays were conducted in 6.5mm 8µm pore polycarbonate membrane Transwell plates (Corning, 3422) coated with 50 *μ*g mL^-1^ rat-tail collagen type I for 1 h at 37 °C and rinsed in PBS before use. Cells were serum starved for 18 hours in enriched DMEM/F12 lacking EGF and Horse Serum prior to seeding at 50,000 cells per well in the serum free media. Transwell migration occurred over 20 hours with the bottom chamber filled with complete Enriched DMEM/F12. Samples were fixed, stained with DAPI, imaged, and the number of migrated cells was counted for each field of view.

### Hanging drop spheroid formation

Spheroids were formed with uninduced cells over 24 hours with 5,000 or 10,000 cells per spheroid in 25 µL hanging drops containing type 1 collagen (20 µg mL^−1^) and 10% [v/v] of 1% [w/v] 25 cP methyl cellulose (Sigma-Aldrich M6385). Co-culture spheroids were generated by combining hanging drop solution of MCF10A cells and ASCs in a 1:1 ratio. Spheroids were rinsed with PBS and media to remove spheroid formation solution before transfer to suspension culture or collagen embedding. For experiments requiring CellTracker (ThermoFisher Scientific C34551) stained spheroids, spheroids were stained in suspension before embedding in serum free media on a rocker at a concentration of 1:2000 CellTracker (5µM) for 30-45 minutes. Spheroids were rinsed in full growth media before collagen embedding.

### Collagen embedding

Spheroids were collagen embedded into custom made PDMS imaging wells that were surface functionalized with 1% [v/v] polyethylenimine (Sigma-Aldrich P3143) and 0.1% [v/v] glutaraldehyde (ThermoFisher Scientific ICN19859580) as previously described^32^. In agreement with previously established protocols^32^, rat tail collagen I (Corning) was diluted to 6 or 2.5 mg mL−1, neutralized with sodium hydroxide (NaOH), and 10% [v/v] 10X DMEM/F12 was used as a phenol red pH indicator. Gels were cold cast over 3 incubation periods of 15 minutes at increasing temperatures of 4 °C, 20 °C, and 37 °C to promote fiber formation. Plates were continually flipped during polymerization to prevent spheroids from sinking to the bottom of the wells. Doxycycline containing induction media was added to each well after polymerization (unless otherwise specified as uninduced) and cells were cultured for 2-4 days. Spheroids were brightfield imaged on the day of embedding and on subsequent days of culture before fixation. For co-embedding studies where ASCs were in the bulk collagen surrounding MCF10A monoculture spheroids, ASCs were added to the collagen solution during collagen dilution.

### Suspension culture

Formed spheroids were transferred into individual wells of 12 well plates coated with 1.5% agarose diluted in DMEM/F12. Plates were cultured at 60 rpm an orbital shaker at 37°C and 5% CO_2_. Spheroid size was monitored by brightfield imaging during the 2-4 day culture period.

### 3D culture in Matrigel

Spheroids were embedded in 70% [v/v] Matrigel (Corning, 356237) in custom PDMS microwells. Polymerization occurred at 37°C with the plate flipped every 5 minutes to prevent spheroid settling.

### Fixing and staining

Samples were fixed for 15 minutes (2D cultures) or 25-30 minutes (3D cultures) in 4% PFA (Electron Microscopy Sciences, 19208) at room temperature. Fixed samples were rinsed and stored in PBS at 4°C before staining or imaging. For IF stained spheroids, samples were permeabilized in 0.1-1% PBS-X for 15 min, blocked in 1% BSA for 1 hr, and stained with AlexaFluor-488, 568, or 647 phalloidin (ThermoFisher, A12379, A12380, A22287) and DAPI (ThermoFisher #D1306) for 1.5 hours at room temperature. Sample stained for α-SMA (Abcam, ab124964) or Collagen IV (Abcam, ab6586) were incubated overnight in 1% BSA and 0.05% PBS-X at 4°C after the blocking step above. The next day, after 3 rinses in PBS, the corresponding secondary antibody was then included in the phalloidin step described above. Samples were rinsed 3x in PBS and stored in PBS at 4°C. Staining for HA was performed using Biotinylated Hyaluronic Acid Binding Protein (Sigma 385911, 10µg/mL) in combination with AlexaFluor 568 conjugated streptavidin (ThermoFisher S11226, 10µg/mL).

### Histology

The Cornell College of Veterinary Medicine Animal Health Diagnostic Center paraffin embedded, sectioned, and H&E-stained collagen embedded and suspended spheroids. Unstained sections were used for HABP and DAPI staining. Slides and tissue sections were imaged on the Aperio ScanScope CS2 (Leica) utilizing a 20x and 40x objective.

### Brightfield, Epifluorescence, and Confocal Imaging

Brightfield, phase contrast, and epifluorescence imaging was done using a Zeiss Axio Observer inverted microscope using the following objectives: EC Plan-Neofluar 2.5X/0.075, EC Plan Neofluar 10X/0.3, EC Plan-Neofluar 20X/0.5, EC Plan-Neofluar 20X/0.4 Ph2 Korr. Confocal images were acquired using a Zeiss LSM 710 and inverted 880 microscopes. Objectives used included C-Apochromat 10x/0.45 W, EC-Plan Neofluar 10X/0.3 A, LCI Plan-Apochromat 25x/0.8 Imm Korr DIC M27 and C-Achroplan 32×/0.85 W Korr M27. Collagen fiber structure was observed with label-free confocal reflectance imaging. Images were acquired at 512×512 pixels with 4 times averaging. Acquired images of spheroids were Z-stacks and tiled as required by the sample size. Images are brightness/contrast-adjusted across conditions for display purposes only while all quantification and analysis was done on unadjusted images.

### Optical Coherence Microscopy and Image Reconstruction

To perform volumetric OCM imaging, the fixed, hydrogel-embedded spheroids were placed on a coverslip or imaged directly in the culture vessel (glass bottom petri dish). OCM imaging was performed with a custom-built, fiber-based, spectral domain OCM setup operating at a center wavelength of 1300 nm^46^. The sample arm utilized a 20x objective with an effective numerical aperture of 0.4. The system axial resolution in tissue (assuming refractive index of 1.33) was 5.6 um and the transverse resolution was 1.6µm. A 1x 1 mm field of view was acquired with 1024×1024 pixels. The imaging range in tissue was 2.6 mm. Acquisitions were made with 50µs exposure time and a frame rate of 15 Hz. The acquired raw data was processed using into complex OCM volumes using background subtraction, linear in wavenumber interpolation, dispersion compensation, and Fourier transformation. Additionally, coherence curvature correction, tilt and computational defocus correction were performed to obtain the final OCM volumes^45^. Lumens within spheroids were segmented using ilastik^101^, with additional manual corrections and smoothing performed in MATLAB. The three-dimensional reconstructions (Fig. 1e) were performed in Dragonfly (Comet Group).

### Image analysis

Images were analyzed using FIJI (ImageJ) and MATLAB. Percent change in spheroid size over time was calculated as ((end size – initial size)/ initial size)*100% using measurement inputs from manual spheroid traces generated from brightfield images in FIJI. Collagen intensity analysis was conducted through the generation of rectangular regions of interest, averaging the reflectance intensity as function of distance from the spheroid-matrix interface, normalizing the values to the average intensity in a distant region of collagen, and setting the average bulk collagen intensity of each condition to a normalized value of 1 set from the average intensity 25-200µm away to account for variance in imaging depth between samples. Nuclear orientation analysis was conducted via a custom FIJI macro built on nuclear ROIs generated via StarDist^102^ whereby isolated nuclear ROIs were further color coded by their aspect ratio, and major axes were isolated and plotted from the shape data. Polyacrylamide bead deformation was quantified through aspect ratio measurements of thresholded beads obtained from the “analyze particles” and “shape descriptor” tool in FIJI. Cell shape from H&E spheroid sections were generated via manual cell tracing and shape quantification in FIJI. In confocal images of the co-culture experiments, radial distance crossings were computed using both FIJI and MATLAB with custom code^81^. Briefly, regions of interest of spheroid invasion were generated in FIJI through thresholding F-actin-stained spheroid and using the analyze particles tool to generate spheroid ROIs. Only collectively invading cells were included in the ROIs with single invading cells excluded from the ROI. These ROIs were then imported into MATLAB for further processing. From the imported ROIs, the centroid of the shape was computed and the Euclidean distance from each perimeter point to the centroid was calculated. The average radial distance was plotted on top of a scatter plot of the moving average-smoothed radial distance from each perimeter point. The number of crossings between each line was recorded along with the standard deviation of the radial distances in each sample.

### Ablation

Spheroids used for ablation experiments and multiphoton imaging were stained in suspension and ablated after or on day 2 of culture in the collagen hydrogels. Multiphoton imaging of cells during and acutely following the production of ablated tracks was performed on a custom-built imaging microscope with a Ti:Sapphire laser tuned to either 880-nm (MIRA, Coherent), or 900-nm (Chameleon, Coherent) as the imaging excitation source. Cells were labeled with CellTracker Orange (CMRA, ThermoFisher) and prior to imaging, growth media was removed and replaced with serum-free, phenol red-free DMEM/F12 supplemented with fluorescein isothiocyanate (FITC) (10 µM, Sigma) to contrast the hydrogel media with produced cavitation bubbles. The collagen fibers within the matrix were visualized by the second-harmonic generation they produce at 440-nm or 450-nm emission for 880-nm and 900-nm excitation, respectively. Imaging and ablation were performed through a 20x, 0.95 NA objective (XLUMPLFLN20XW, Olympus). The detection optics consisted of a 735-nm primary long-pass dichroic mirror, 488-nm secondary long-pass dichroic and 560-nm tertiary long-pass dichroic (AVR-Optics) with additional bandpass filters at 439/154 nm (and later 447/60 nm) for imaging of SHG, 510/42 nm for imaging of FITC, and 617/73 nm for imaging of CellTracker Orange (AVR-Optics) placed in front of the respective photomultiplier tube to separate out the emission from each component. Spheroid samples were placed upon a Newport XPS Series 3-axis stage for producing the tracks. ∼50 femtosecond laser pulses at 1 kHz from a chirped pulse amplifier system^103,104^ (Legend, Coherent) were centered within the field of view while the sample was translated at 15 µm/s using the Newport stage. A custom program in MATLAB (Mathworks) controlled the stage movement and communicated with a microcontroller (Arduino Uno) to gate a mirrored shutter (LS6 ZM2-100, Uniblitz) such that the amplifier pulses ablated the media only during the intended creation of the tracks. Pulse energies were adjusted between 1-3 µJ using an ND-filter wheel in conjunction with a half-wave plate (AHWP05M-980, ThorLabs) and polarizing beamsplitter cube (PBS252, ThorLabs) so that the produced tracks were 30-80 µm in width and did not produce uneven blebbing along the track. The cell tracks were 200 µm in length and consisted of two layers spaced 5 µm apart in depth. Tracks were placed along edges of the spheroid with relatively smooth walls and without indents or folded collagen layers. A total of two tracks were placed on opposite sides of each of the embedded spheroids, as far apart as possible based on the prior criteria. Three-dimensional image stacks of the top-halves of the spheroids were collected prior to creation of the ablated tracks and 3-5 minutes following. Videos were taken at 3-4 frames per second for 3-5 minutes during and immediately after laser ablation to capture acute flow dynamics following rupture of the collagen wall.

### Analysis of Ablated Spheroids and Calculations of Intraspheroidal Pressure

Particle tracking for ablation experiments was performed with MTrackJ plugin in FIJI. FITC media tracking and cavitation bubble displacement was quantified as change in distance from the plane of the spheroid matrix interface at the ablated path to the edge of the bubble. In this analysis t=0 was set at the point after the stage motion stopped following ablation and change in cavitation bubble displacement was calculated after 120 seconds with manual line segment measurement in FIJI. The Hagen-Poiseuille equation^47,48^ was used to calculate the pressure difference across the spheroid interface with the following parameters manually collected from spheroid videos where particles could be tracked through the ablated pore to model as flow through a pipe in the region of the ablated pore: L = length of the particle movement through the pore, R= radius of the ablated pore, µ = 16.8cp (estimated value based on 1mg/mL 2MDa HA^105^), Q= volumetric flow rate calculated from measured particle velocity multiplied by the area of circle of radius R. Length and radius measurements were computed for each ablated pore to account for heterogeneous pore/channel size.

### Assays and inhibitor studies

Secreted hyaluronan was quantified using the Hyaluronan DuoSet ELISA (R&D systems DY3614). Intraspheroidal HA was released through spheroid lysis via 10 sonication pulses into the cell culture media. Three spheroids were pooled for each sample analyzed via ELISA. Batimastat (Tocris 2961) was used at a concentration of 10µM in enriched DMEM/F12 and replenished every 2 days with media changes. Blebbistatin (Sigma B0560) was used at 20 µM and delivered on the same dosing schedule. The QuantiFluor (Promega E2670) DNA assay was used according to manufacturer instructions. For collagen embedded samples, samples were digested out of collagen hydrogels through incubation with Collagenase type 1 (Worthington Biochemical Corporation LS004197) at 3mg/ml in serum free DMEM/F12 for 1 hour at 37°C. Samples were centrifuged and resuspended in Caron’s Buffer (25 mM Tris-HCl, 0.4 M NaCl, 0.5 % SDS) as pooled samples with 3 spheroids per 25µl of solution. Pooled samples were sonicated ten times to prepare for the assay.

### Polyacrylamide pressure sensors

Deformable polyacrylamide pressure sensor beads^55,56,106^ were obtained from the Max Planck Institute for the Science of Light in collaboration with Jochen Guck. The beads contain AlexaFluor 488 and were characterized to have an average diameter of 12.7 µm and a stiffness on the order of 1 kPa. Beads were incorporated into the spheroid formation solution and into the collagen precursor solution at an estimated density of 2000 beads per mL. Spheroids were cultured for 2 days, fixed, and imaged for bead compression analysis detected via deformation^53^.

### Statistics

At least three biological replicates were used for each experiment unless otherwise noted. Exact sample sizes are listed in each figure legend. Statistical tests were conducted in GraphPad Prism (v.10.4.2) with statistical significance defined as a p-value of less than 0.05. Data is plotted as means with standard deviation error bars unless otherwise indicated. Comparisons of 2 groups were performed with unpaired t-test with Welch’s correction. Data of 3 or more groups with multiple time points or conditions were evaluated using ordinary one-way ANOVA with Sidak’s multiple comparison. Any deviations from the above statistical tests are described in the figure legends.

### Computational Modeling of Matrix Densification and Lumen Formation

Supplementary Note 1 contains the detailed methodology of the Finite Element Analysis used to simulate material deformation in the presence of an expanding sphere. Supplementary Note 2 includes all relevant details, equations, and links to code for our custom simulation of lumenogenesis.

### Code Availability

Cell-vertex modeling Mathematica notebooks are published in Zenodo DOI: 10.5281/zenodo.24411407

## Acknowledgments

Thank you to the entire Fischbach lab at Cornell University for providing feedback on all stages of this research and manuscript preparation. We would like to thank the Paszek Lab: Joe Kuo, Monet Roberts, and Matthew Paszek for generating the MCF10A HAS overexpressing cell lines and sharing these cells with us. Thank you to our collaborators at the Max Planck Institute for the Science of Light; Ruchi Goswami and Jochen Guck for supplying us with the deformable polyacrylamide pressure sensors. Thank you to Lu Ling for her guidance in spheroid assay optimization and imaging advice. We would like to thank the staff of the Cornell Biotechnology Resource Center especially Rebecca Williams and Johanna Dela Cruz for their assistance with instrumentation setup for confocal imaging. Meg McFetridge conducted OCM imaging and we thank her for her contributions. Thank you to Larry Bonassar for the discussions surrounding the calculations of intraspheroidal pressure. We thank the Cornell Histology Core for paraffin embedding, sectioning, and H&E staining spheroid samples. We acknowledge financial support from NCI R01CA259195, R01CA276392, R01CA297166, and the Breast Cancer Alliance. Confocal imaging was conducted at the Cornell Biotechnology Resource Center Imaging Facility (RRID:SCR_021741) with the following funding: NIH S10RR025502 for the Zeiss LSM 710 confocal microscope and NIH S10OD018516 for the Zeiss LSM880 confocal microscope. S.J. and N.B. were supported through NSF-SNSF collaborative program by the U.S. National Science Foundation under Award No. 2516642.

## Notes

### Competing Interest Statement

The authors have declared no competing interest.

