## Extended Data Figs 1-11 for "Cyst-like intraluminal hyaluronic acid accumulation inhibits breast epithelial cell invasion via pressure-driven ECM compaction"

**Extended Data Figures 1-11**

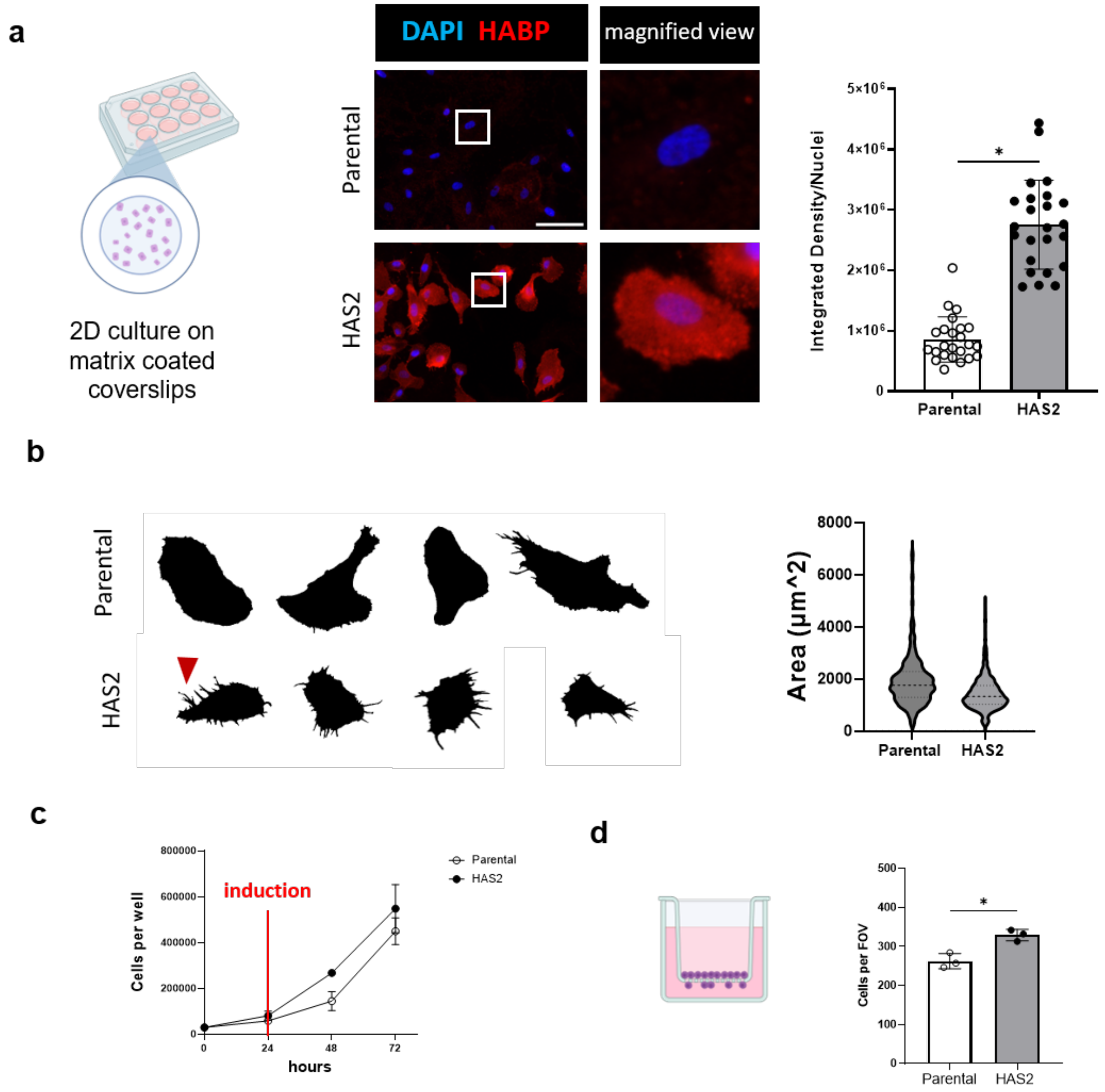


**Extended Data Figure 1: HAS2 overexpressing MCF10As retain cell surface associated HA and display characteristic glycocalyx-associated changes to morphology and invasion while maintaining a comparable proliferation rate**. **a**) Schematic of 2D culture systems displaying matrix coated glass coverslips. Hyaluronic Acid Binding Protein (HABP, red) staining of 2D cultured cells reveals cell-surface associated HA in the HAS2 cells. * indicates statistical significance as determined by a t-test with Welch's correction, n=24 fields of view per condition. **b**) Representative cell masks of F-actin-stained cells on matrix coated coverslips reveal membrane protrusions on the HAS2 cells (red arrow) and a less spread morphology as seen in the plot of cell area. **c**) Manual cell counts of induced cells harvested from 6 well plates. To allow for plate adhesion, cells were induced with doxycycline to overexpress HAS after 24 hours (red vertical line) and counted at 24, 48, and 72 hours. **d**) Plots of a transwell migration assay show the number of cells per field of view that migrated to the bottom of the transwell after 20 hours. Each dot is the average of 6 fields of view for 3 wells. * Indicates statistical significance as determined by a t-test with Welch's correction. Schematic created in Biorender.

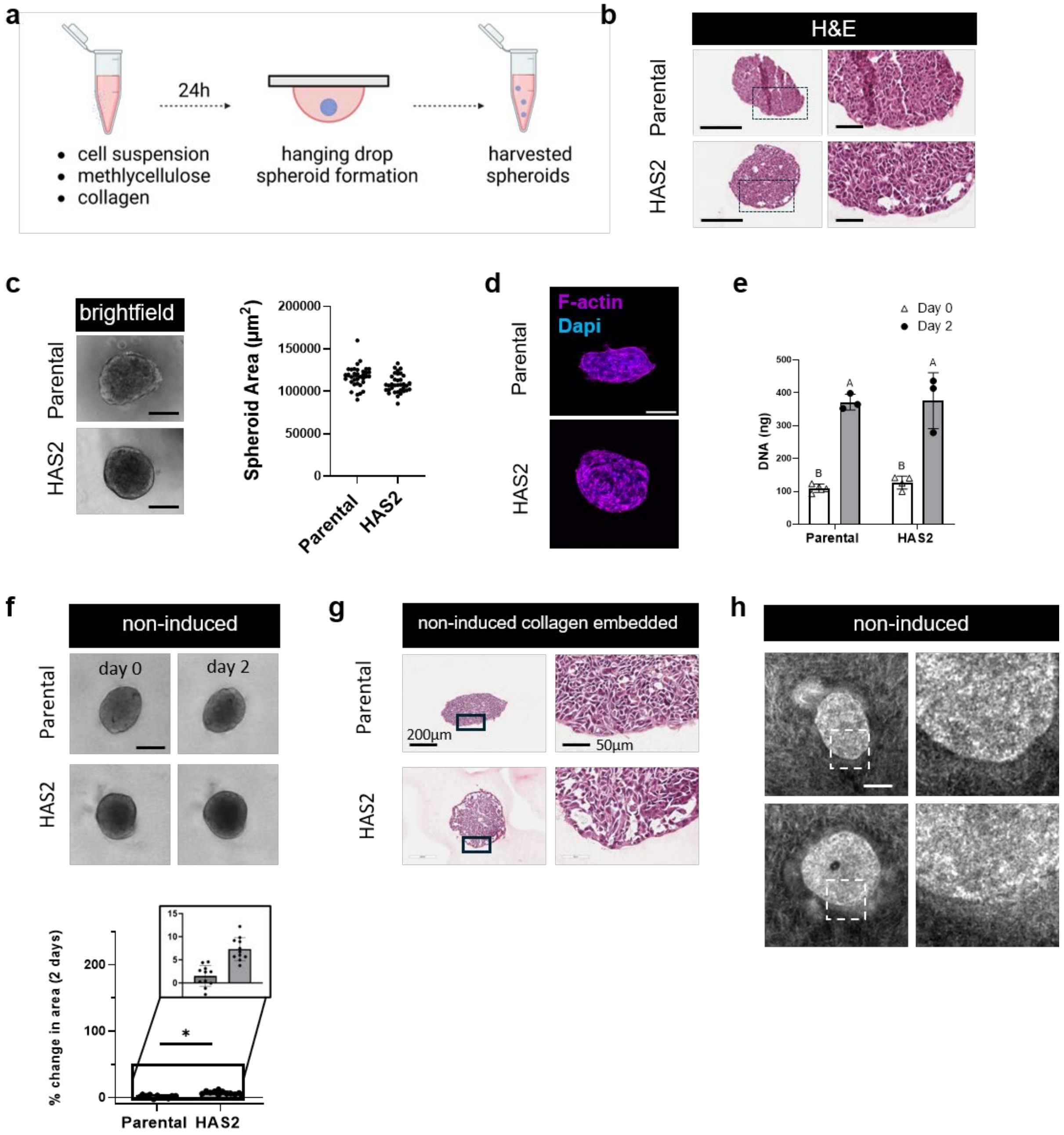


**Extended Data Figure 2: Expanding induced spheroids do not have an increase in DNA content as compared to parental controls, but both expansion and lumen formation are dependent on doxycycline induction of HAS2. a)** Illustration of hanging drop spheroid formation and harvest. Created in Biorender. **b)** Hematoxylin- and eosin-stained sections of fixed spheroids immediately post-harvest from 24-hours of hanging drop formation show the hanging drop method promotes aggregation, cell-cell adhesions, and the presence of only minor cell-free gaps in the spheroid bulk. Scale bar, 200 µm and 50 µm on zoom insets. **c)** Brightfield images of spheroids formed with the hanging drop method and associated quantification of produced spheroid size. Scale bar, 200 µm. **d)** Maximum projection confocal images of day 2 spheroids cultured in 6mg/mL collagen stained with phalloidin (purple) and DAPI (blue) show a lack of invasion. Scale bar, 200 µm. **e)** DNA content of spheroids on day 0 and after 2 days in culture in 6 mg/mL collagen gels. Each data point represents 3 pooled spheroids. n= 3–4 per condition, $p<0.05$. Compact letter display of statistical significance reveals a significant increase in DNA content from day 0 to day 2 and no statistically significant difference between parental and HAS2 spheroids on either timepoint. **f-h** feature collagen embedded spheroids that were not induced with doxycycline to activate the overexpression of HAS2. **f**) Brightfield images and associated size quantification of non-induced 6mg/mL collagen-embedded spheroids. Scale bar =200µm, n=11; *p <0.05. **g)** H&E images of paraffin embedded sections of non-induced spheroids cultured in 6 mg/mL collagen gels for 2 days. Scale bar = 200µm in the top row and 50µm in the magnified insets displayed to illustrate spheroid morphology and lack of invasion at the spheroid-matrix interface. **h**) Optical coherence microscopy images of XY views of day 2 collagen embedded spheroids without doxycycline induction show lack of invasion and lack of peripheral lumen (cell-free region) formation. Scale bar, 200 µm and 50 µm (insets).

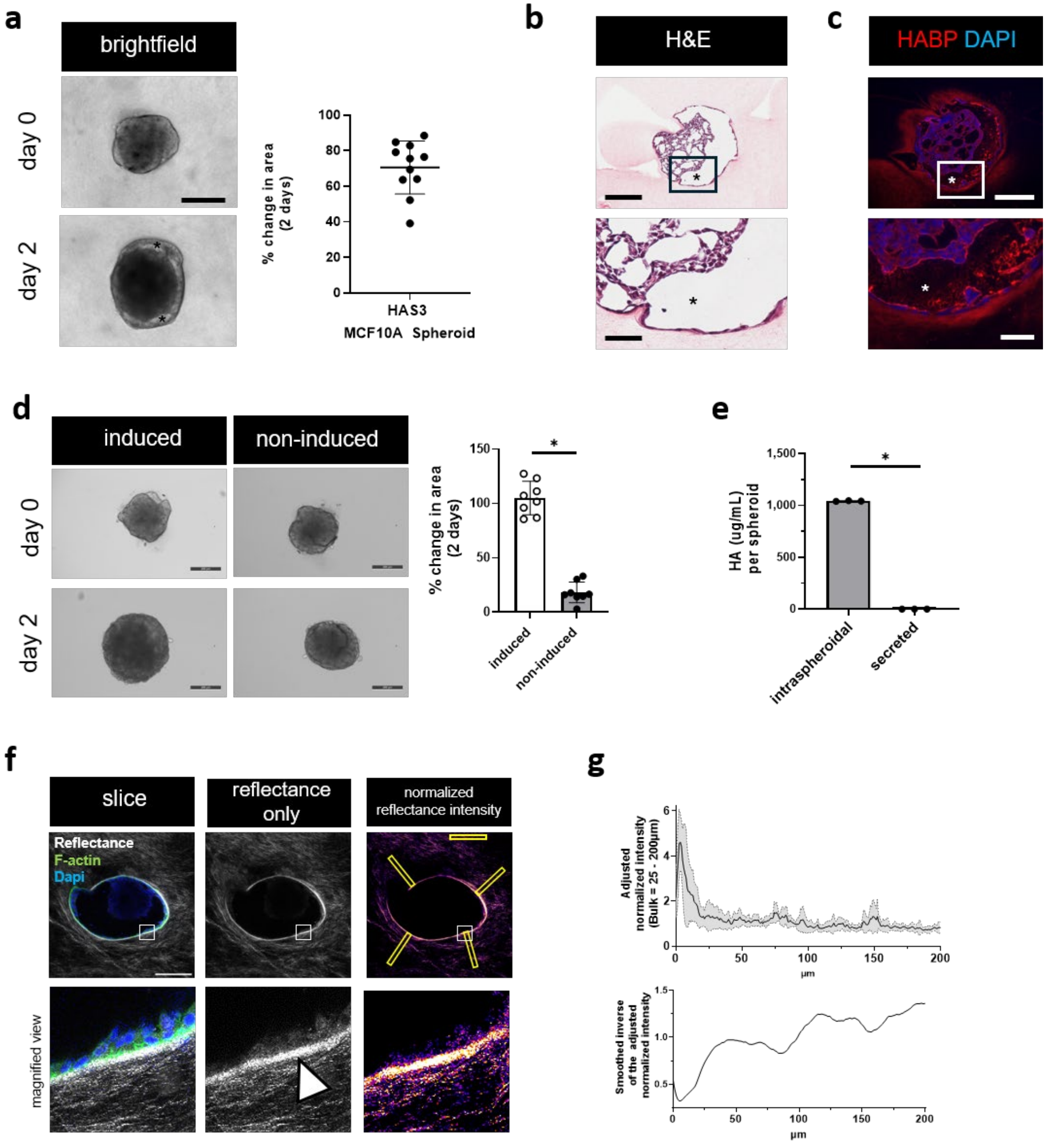


**Extended Data Figure 3: HAS3 overexpressing spheroids display the same expansion phenotype as HAS2 spheroids indicating an HAS isoform-independent phenotype**. **a)** Brightfield images and corresponding quantification of HAS3 spheroids in 6 mg/mL collagen hydrogels. Scale bar=100µm. In a-c * indicates translucent cell-free regions / lumen-like structures **b)** H&E-stained sections of paraffin embedded spheroid in a 6mg/mL collagen hydrogel. Scale bar = 200µm in the overview image and 50µm in the magnified view. **c)** HABP stained paraffin embedded spheroid section with HA positive regions shown in red and DAPI displayed in blue. Scale bar = 200µm in the overview image and 50µm in the magnified view. **d)** Brightfield images of suspension cultured spheroids and corresponding quantification of percent change in area show expansion is an induction dependent phenotype. Scale bars =200µm. **e)** HA ELISA results of an analysis of lysed spheroids show orders of magnitude higher concentrations of HA accumulated inside the spheroid as opposed to secreted into the surrounding cell culture media. Each data point is 3 pooled spheroids. **f)** Confocal images of a HAS3 collagen embedded spheroid with F-actin in green, DAPI in blue, and collagen reflectance in grey. White arrow highlights the densified collagen. Yellow boxes on the normalized intensity display (rightmost image set) indicate regions of interest for the background and 4 radial samples of collagen used for intensity measurement. **g)** Top plot is the average ± SD of the normalized collagen intensity collected from the 4 measured radial regions of interest starting from the spheroid surface and extending 200µm into the matrix. Bottom plot is the smoothed inverse of the average normalized intensities.

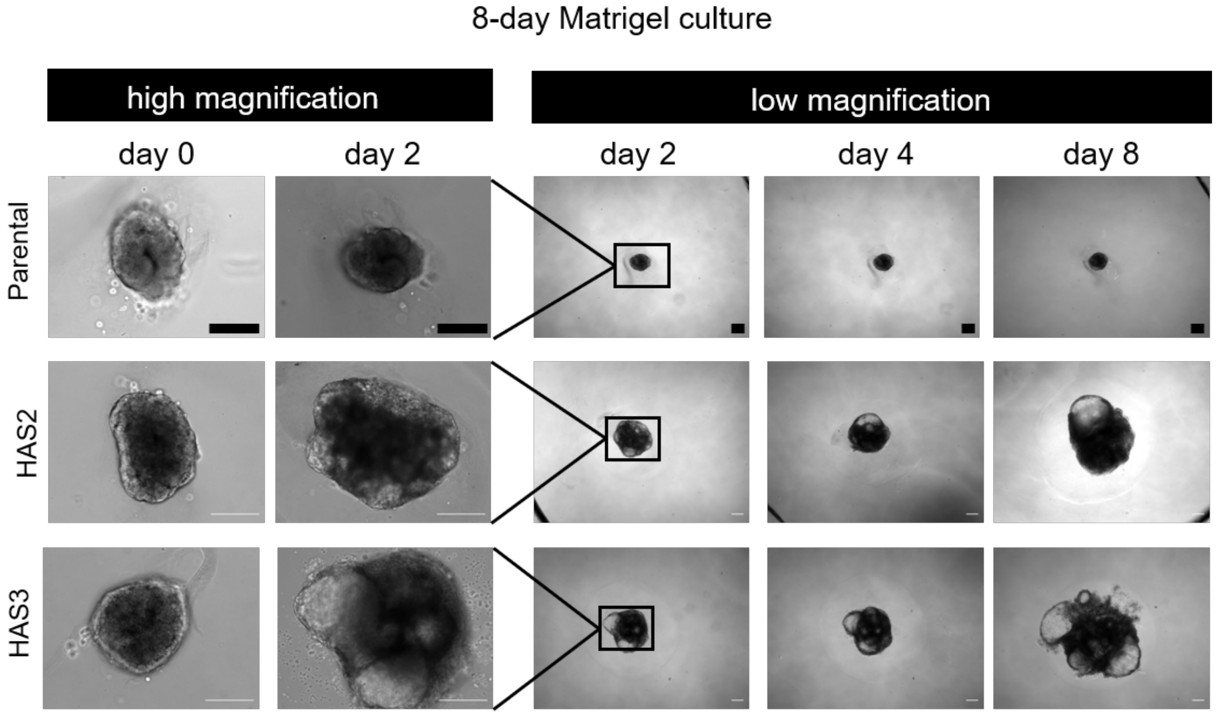


**Extended Data Figure 4: HAS2 spheroids expand in Matrigel.** Brightfield images on days 0 and 2 were taken with a 10X objective. Day 2-8 images taken with a 2.5X objective. All scale bars are 200 µm.

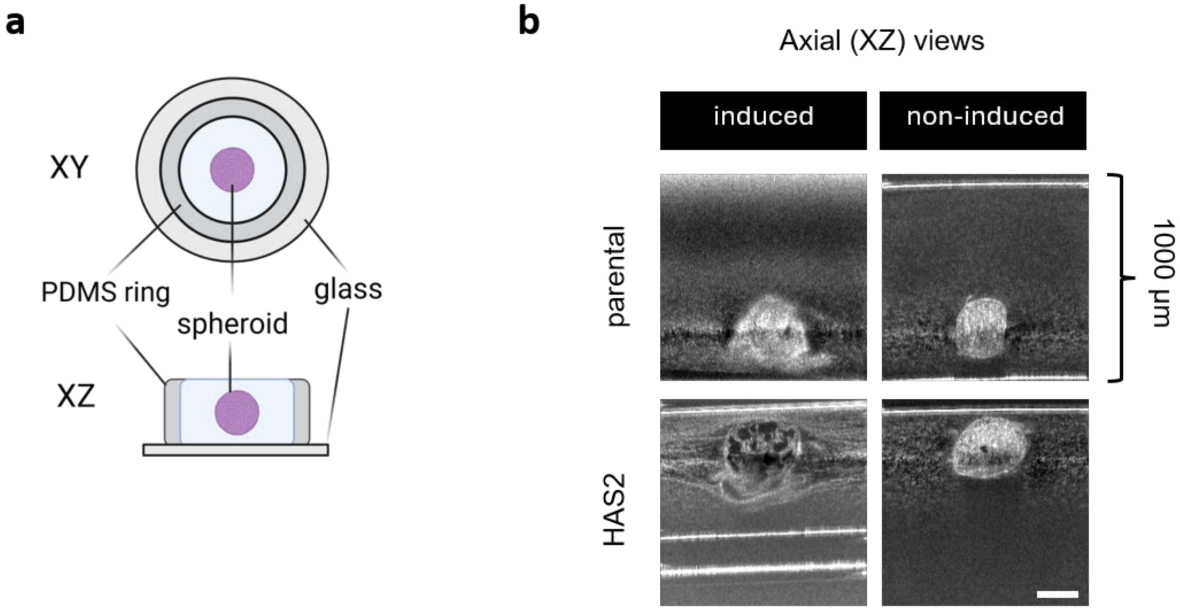


**Extended Data Figure 5: OCM Axial (XZ) view spheroid samples presented in Figure 2**. a) Schematic of the imaging planes of the spheroids in the PDMS imaging chambers. b) XZ views of the spheroids displayed in Fig. 1 with a field of view of 1mm in the X direction.

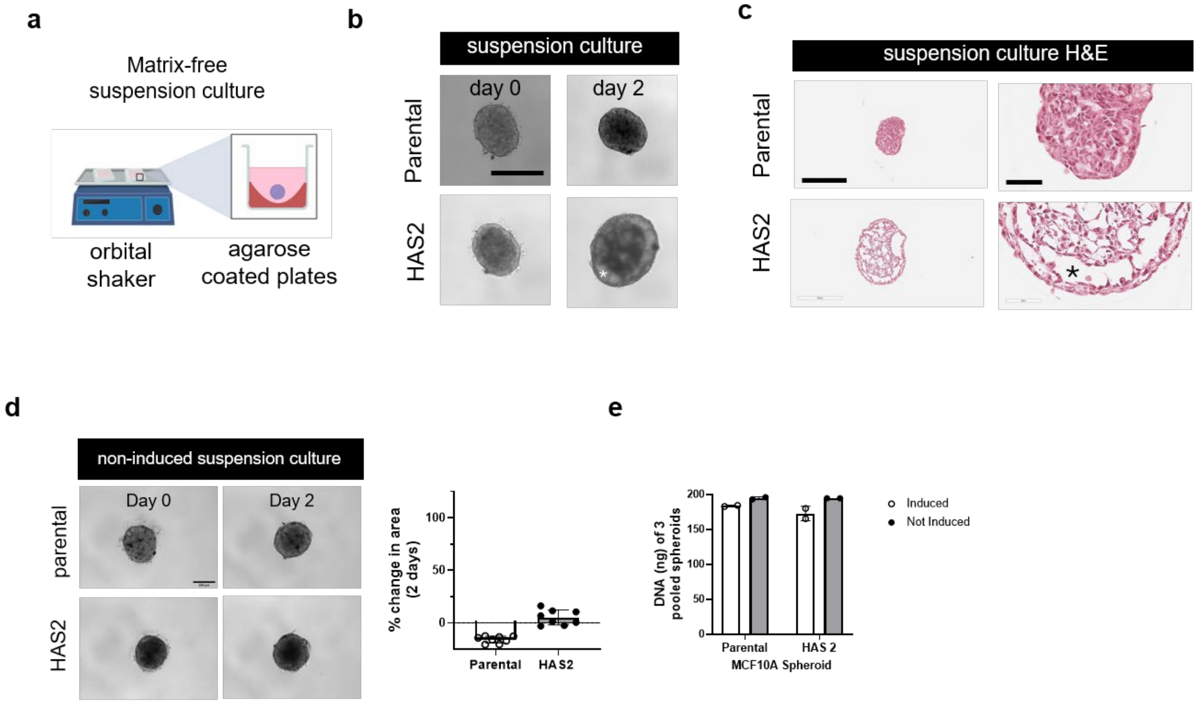


**Extended Data Figure 6: Suspension culture spheroids form lumens when induced but display altered morphology in the outer cell layer when compared to spheroids confined by a collagen hydrogel. a)** Schematic of suspension culture of spheroids in agarose coated plates on an orbital shaker. Created in Biorender. **b)** Brightfield images of suspension culture spheroids. * shows translucent region. Scale bar =200µm. **c)** H&E images of suspension culture spheroids on day 2 of culture. * indicates cell-free space, scale bar = 300µm and 60µm in the magnified views (right image in each row). **d)** Brightfield images of suspension culture spheroids cultured for 2 days in the absence of doxycycline with corresponding quantification of percent change in area. Scale bar =200µm. **e)** DNA assay of suspension culture spheroids with and without doxycycline induction. Each data point is 3 pooled spheroids.

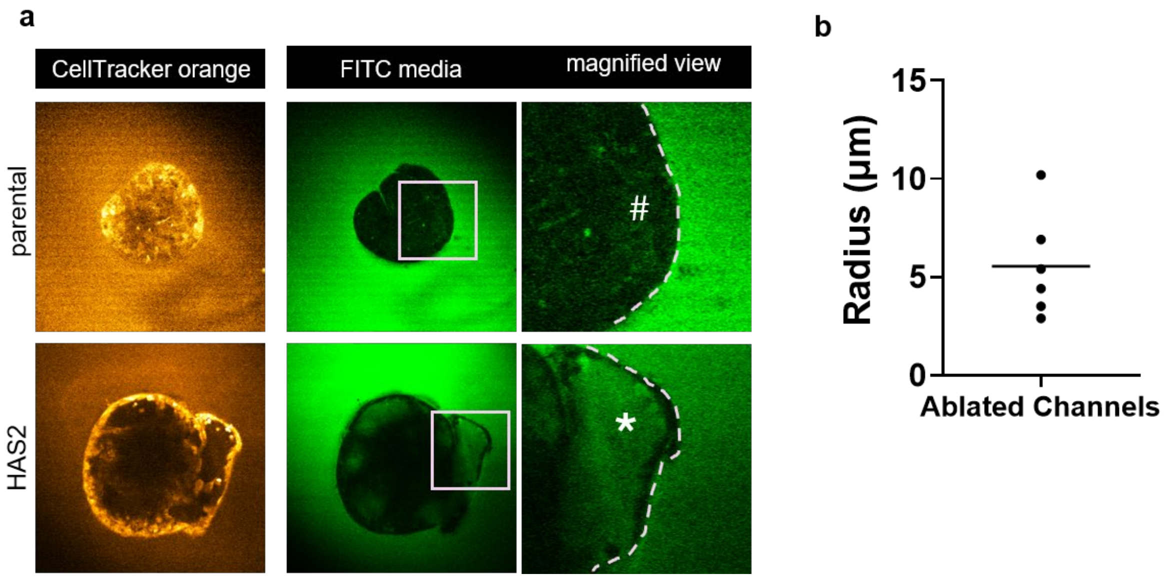


**Extended Data Figure 7: Pre-ablation images of collagen embedded CellTracker orange-stained spheroids in FITC containing cell culture media and quantification of ablated microtrack radii.** **a)** CellTracker Orange staining of cells (orange) and FITC staining of cell culture media and intraspheroidal fluid (green). The white dotted lines in the rightmost magnified insets demarcate the spheroid-matrix interface. # indicates a dense cellular region in the parental spheroid that is devoid of interpenetrating FITC labeled fluid. *indicates a cell-free lumen region in the HAS2 spheroid filled with FITC labeled fluid. Images histogram normalized in FIJI for display consistency. **b)** Quantified radius of ablated channels measured as the exit site of the spheroid (the gap in the outermost cell layer, not the full width of track located distally from the spheroid-matrix interface). Each data point is one ablated channel.

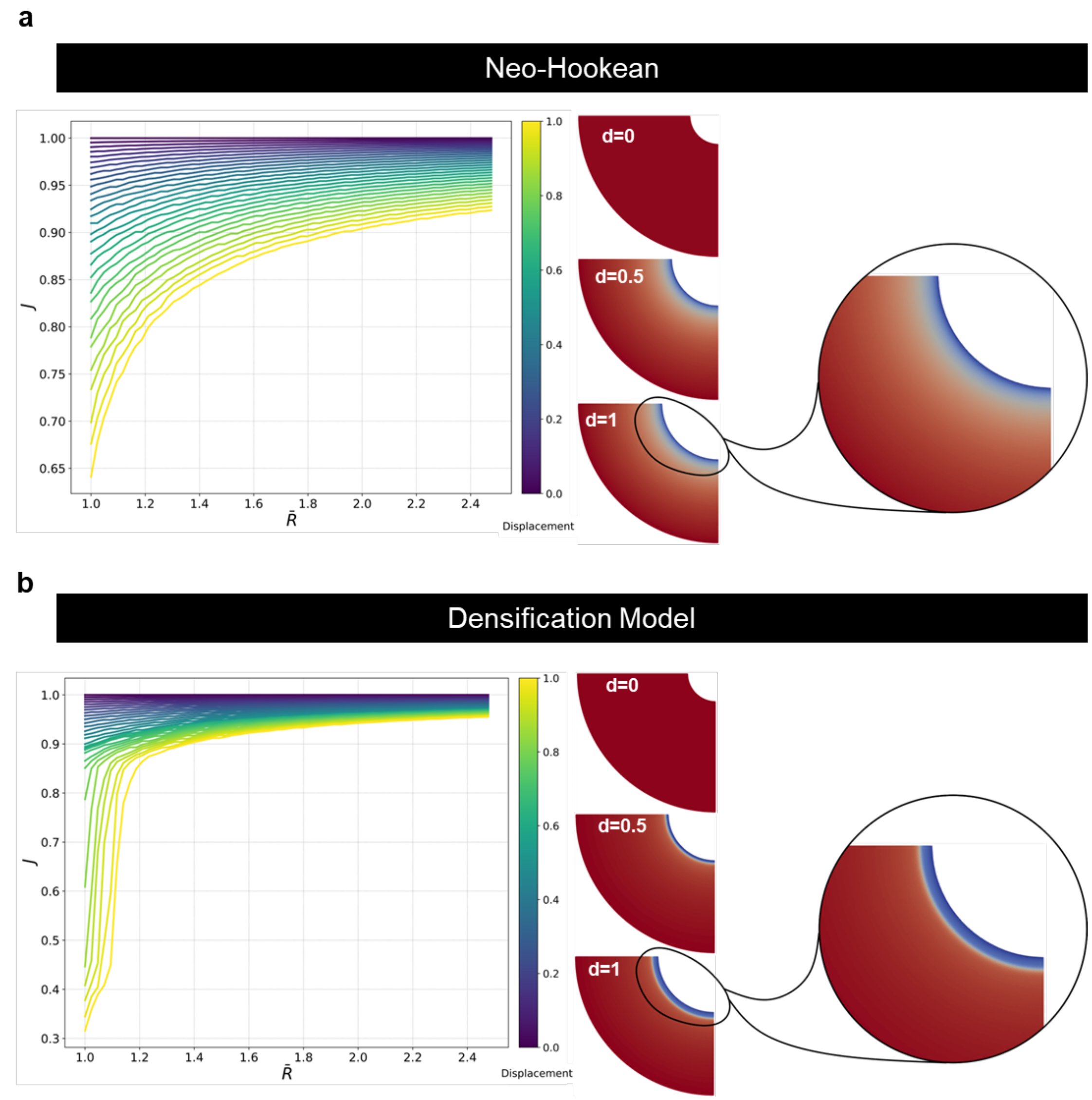


**Extended Data Figure 8: Time course pressurized cavity expansion of a compressible foam.** Line plots of Jacobian (volume ratio) with respect to the radial position, as pressure is applied over a series of 30 load steps to recapitulate spheroid expansion over time. Matrix quadrants at displacement=0,0.5, and 1, spatially display the Jacobian values as a function of radial distance on a blue to red color gradient. **a)** In the Neo-Hookean model there is no indication of sharp transitions and corresponding phase transitions. **b**) the densification model shows the emergence of steep slopes in the Jacobian plot as a function of time.

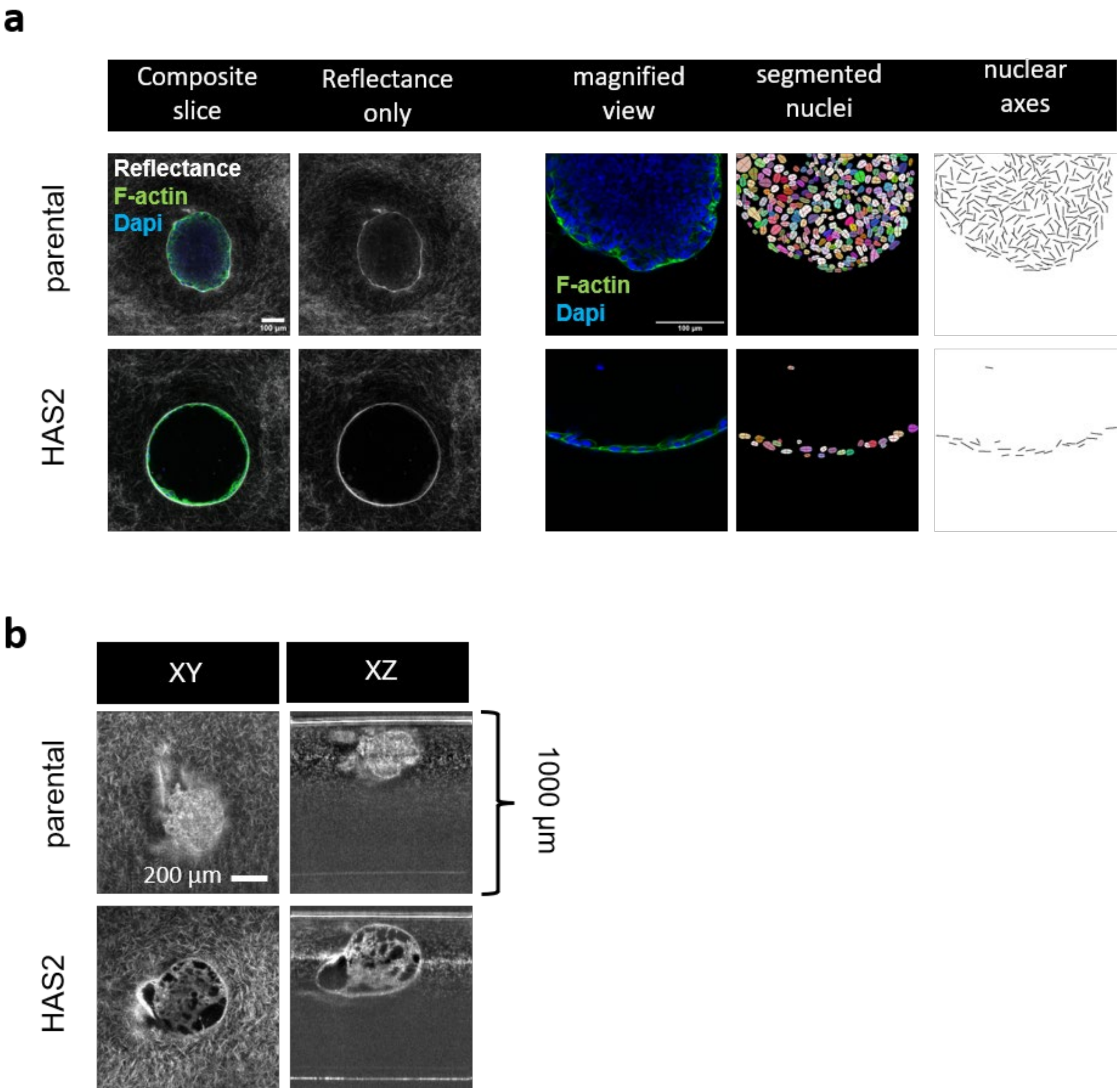


**Extended Data Figure 9: Characterization of spheroids embedded into 2.5 mg/mL collagen hydrogels**. **a**) Confocal imaging of DAPI (blue) and F-actin (green) stained spheroids with collagen reflectance displayed in greyscale. Scale bar = 100µm in the overview images (left panel) and magnified views (right panel). Nuclear segmentation completed with the StarDist plug-in in FIJI. Nuclear axes were drawn with a custom FIJI script **b**) OCM images of collagen embedded spheroids showing the presence of signal free lumens in the HAS2 spheroid.

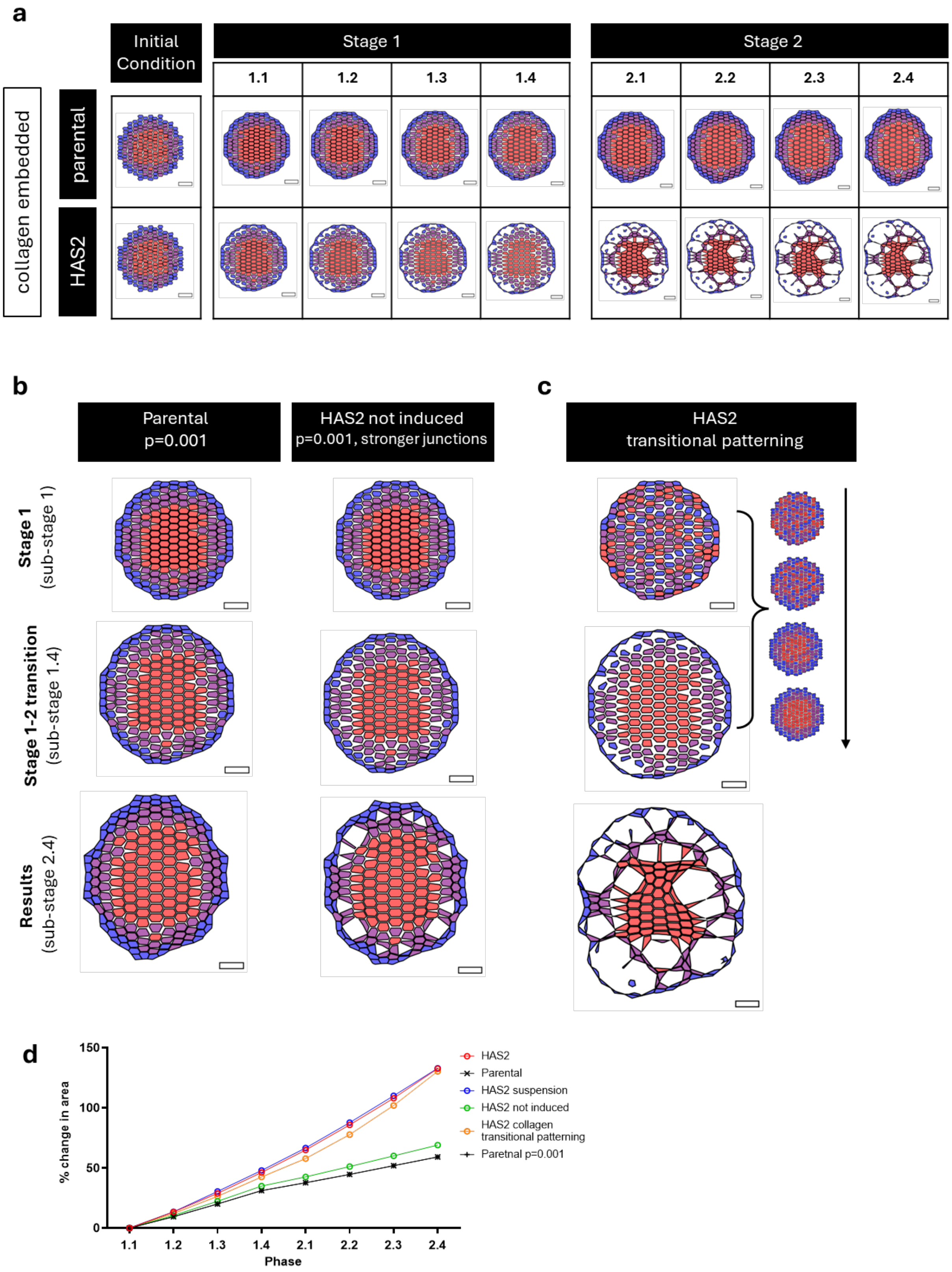

a
Initial Condition
Stage 1
Stage 2
1.1
1.2
1.3
1.4
2.1
2.2
2.3
2.4
collagen embedded
parental
HAS2
b
Parental p=0.001
HAS2 not induced p=0.001, stronger junctions
Stage 1 (sub-stage 1)
Stage 1-2 transition (sub-stage 1.4)
Results (sub-stage 2.4)
c
HAS2 transitional patterning
d
% change in area
Phase
150
100
50
0
HAS2
Parental
HAS2 suspension
HAS2 not induced
HAS2 collagen transitional patterning
Paretnal p=0.001

**Extended Data Figure 10**: **Lumen formation requires junctional breaks. The location of the junctional breaks at the end of Phase 1-4 (not during the rest of phase 1) determines the final intraspheroidal lumen location.** **a**) Time course results of the collagen embedded parental and HAS2 conditions to show all stages of the simulations featured in figure 4. **b**) Here, we modified the parental condition to have the same pressure differential as in the HAS2 conditions but overserved no lumen formation due to the relative higher spring constants used in this condition. To model our experimental condition of the non-induced HAS2 spheroids, we increased the spring constants (to recapitulate the stronger cell-cell junctions in the absence of glycocalyx production) but maintained the pressure differential. The results mirror those seen in non-induced samples in vitro with small peripherally located lumens. **c**) Implementing the sorting of cells in the HAS2 collagen embedded model based on adhesion strength during phase 1 as opposed to at the start of phase 1 results in a very similar bulk morphology as compared to the pre-sorted condition (**Fig. 4**). **d)** Simulated spheroid cross section measurements to quantify the overall percent change in area, or the relative increase in size in the cross section as a function of simulation phase.

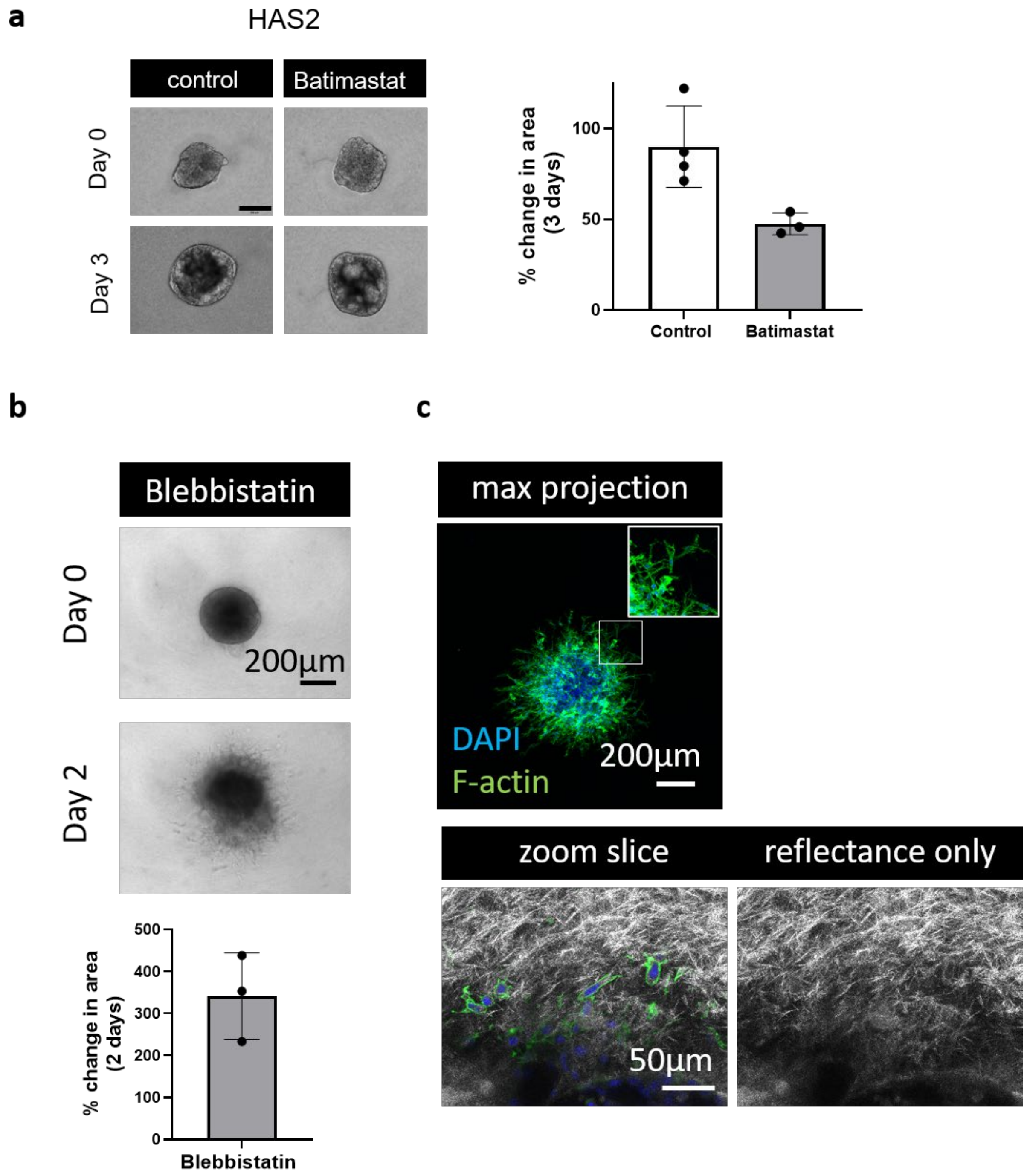


**Extended Data 11: Inhibitor treatments of mono- and co-culture spheroids. a)** Batimastat treatment reduced expansion (% change in area) but did not prevent the formation of translucent lumen-like structures as observed with brightfield imaging. 3-day culture period, scale bar =200µm, each point represents one spheroid. **b-c)** treated co-culture spheroids over 2 days. **b)** Brightfield images and quantification of percent change in area of Blebbistatin treated spheroids show invasion persists despite inhibiting the contractile phenotype of the stromal cells. **c**) Confocal image of a Blebbistatin treated spheroid with F-actin in green, DAPI in blue, and reflectance in grey. No collagen ring is observed at the spheroid matrix interface.
