## Supplementary Note 1 for "Cyst-like intraluminal hyaluronic acid accumulation inhibits breast epithelial cell invasion via pressure-driven ECM compaction"

### Supplementary Note 1. A computational model of spheroidal expansion and collagen densification.

We base the general theory of kinematics of a soft body, focusing on phase transitions that can accommodate densification. Computational models of three-dimensional tissue deformation provide a framework for understanding the mechanical response of biological micro tissues[1]. Specific considerations for implementation of this work are introduced in the following formulations.

#### 1. Kinematics

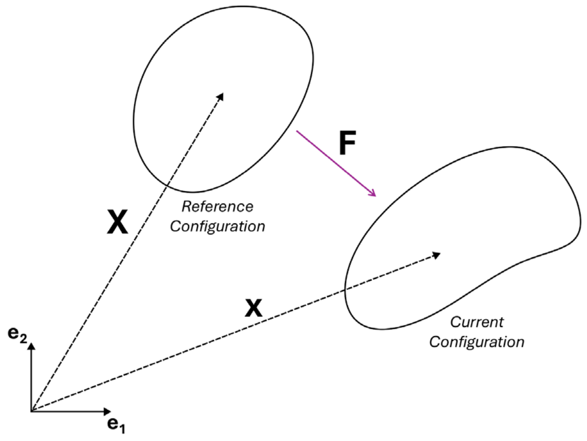


Fig S1: Illustrating the reference and current configuration of the continuum body.

Fig. S1 explicitly establishes the notations for the continuum body, $\mathcal{B}$ in the reference and the current configuration. The frame of reference illustrated is Cartesian coordinate system. The two configurations of $\mathcal{B}$ , are depicted by, $\Omega$ and $\Omega_0$ for current and reference configurations. If the current configuration coincides with the reference configuration, we consider the body to be "stress-free". The position vector for reference and current configuration is denoted by, $X$ and $x$ respectively. We use the notation: $\chi\colon\ V \rightarrow \mathbb{R}^3$for the deformation of body $\mathcal{B}$ . Based on the position vectors, we define the motion $\chi$ as the vector field of the mapping, $x = \chi(X)$. This helps defining the kinematics of a reference point as,

$$u(X,\ t) = x(X,\ t) - X \tag{S1}$$

where, $u(X,\ t)$ describes the displacement vector field. Furthermore, the kinematics of an infinitesimal bulk element are described by,

$$F(X,\ t) = \frac{\partial \chi(X,\ t)}{\partial X} = \nabla_X x(X,\ t) \tag{S2}$$

$$F^{-1}(x,\, t) \;=\; \frac{\partial \chi^{-1}(x,\, t)}{\partial x} = \nabla_x X(x,\, t) \tag{S3}$$

where $F(X,\, t)$ and $F^{-1}(x,\, t)$ are the deformation gradient and inverse deformation gradient, respectively. Note that $J(X,\, t) \;=\; \frac{dv}{dV} = \det \quad F(X,\, t)$ is the Jacobian determinant defining the ratio of a volume element between reference and spatial configuration. Motion of an arbitrary differential vector element can be mapped by the deformation gradient, $F$ , following the standard framework of nonlinear continuum mechanics[2] .

### 2. Helmholtz Free Energy Density

The observed phenomenon of densification of collagen needs to be encapsulated by formulation of Helmholtz free energy function, governing the state of the system. Previous studies have demonstrated the importance of mechanical stresses in governing the morphological response of fibrous micro tissues[3] . We decompose the free energy density function, $\psi$ into local and non-local contributions to account for both intrinsic material properties and spatial interactions.

The local component, $\psi_{loc}$ represents the conventional elastic response of the material, describing its resistance to deformation. While, the non-local component, $\psi_{non-loc}$ incorporates the spatial interactions that arise due to the network architecture encountered in the base material. Thus, Helmholtz free energy density function is expressed as:

$$\psi = \psi_{loc} + \psi_{non-loc}$$

Focusing on the local description of elastomeric material, we denote Helmholtz free energy density function primarily as $\psi_{loc}(F)$. The observed compressive trends lead us to choose compressive Neo-Hookean constitutive model for describing the local component.

$$\psi_{loc}(F) \;=\; \frac{\mu}{2}(I_1 \;-\; 3 \;-\; 2\ln J)$$

Furthermore, non-local description of Helmholtz free energy density function, $\psi_{non-loc}$, needs to capture the essence of phase transition. Phenomenologically, the densified and the rarefied region demands the presence of a double-well potential, leading to formulate the non-local description.

$$\psi_{non-loc} \;=\; 10\kappa(J \;-\; 0.2)^2(J \;-1)^2 + \frac{\kappa}{2}(\ln J)^2$$

Thus, the complete Helmholtz free energy density is expressed as,

$$\psi = \frac{\mu}{2}(I_1 - 3 - 2\ln J) + 10\kappa(J - 0.2)^2(J - 1)^2 + \frac{\kappa}{2}(\ln J)^2$$

### 3. Potential Energy and Weak Form

The spheroidal expansion leading to matrix densification requires computation of total potential energy, An equilibrated configuration is obtained by minimizing this functional considering all admissible deformations. While more general continuum formulations may incorporate additional boundary potential energies and their corresponding finite element treatments[4,5], the present formulation considers the bulk Helmholtz free energy, with spheroidal expansion prescribed through displacement boundary conditions. To determine the finite element solution, we derive the weak form of the governing equations by requiring the first variation of the total potential energy to vanish, $\delta\Pi(\chi) = 0$. The resulting weak form is expressed as,

$$G = \int_{\Omega_0} P : \nabla_X \delta u \, dV - \int_{\Omega_0} b \;\delta u \, dV - \int_{\delta\Omega_0} t \;\delta u \, dS$$

where, $\delta u$ is the admissible deformation field, with $b$ , $t$ being body force and traction applied on the infinitesimal volume and surface respectively. The problem is highly non-linear due to presence of phase transitions. To address this, we introduce a non-local variable[6], $\tilde{J}$, a regularized counterpart to local volume ratio $J$, to follow the volumetric deformation, penalize its gradients, and stabilize the solution when the open-source platform FEniCS[7] is employed for the finite element simulation. We also used the Scalable Nonlinear Equations Solvers (SNES) from the open-source toolkit PETSc[8], which provides numerical computations of a Newton-type iterative procedure to solve the nonlinear variational problem.

### 4. Finite Element Implementation

The reference configuration corresponds to the state where the matrix surrounding the spheroids are in the rarefied regime, setting the initial condition of $\tilde{J} = J = 1$ , while the incremental displacement is applied on the inner boundary, simulating the expansion of spheroids in the matrix. Furthermore, we assume no external mechanical loads and thereby set body force and traction to zero. This helps isolate the internal mechanical response due to the strain energy density function implemented. The refinement in the strain energy density employed in the simulation, captures the experimental observation of densified phase. The incremental expansion of the spheroid establishes a densified region near the expansion, while the matrix away from the cell, maintains its rarefied form.

### Supplementary Note 1 References


1. Kim, J., Mailand, E., Ang, I., Sakar, M. S. & Bouklas, N. A model for 3D deformation and reconstruction of contractile microtissues. *Soft Matter* **17**, 10198–10209 (2021).
2. Holzapfel, A. G. Nonlinear solid mechanics II. (2000).
3. Mailand, E., Li, B., Eyckmans, J., Bouklas, N. & Sakar, M. S. Surface and Bulk Stresses Drive Morphological Changes in Fibrous Microtissues. *Biophys. J.* **117**, 975–986 (2019).
4. Javili, A. & Steinmann, P. A finite element framework for continua with boundary energies. Part II: The three-dimensional case. *Comput. Methods Appl. Mech. Eng.* **199**, 755–765 (2010).
5. Steinmann, P. On boundary potential energies in deformational and configurational mechanics. *J. Mech. Phys. Solids* **56**, 772–800 (2008).
6. Joshi, S. *et al.* Instabilities and phase transitions in architected metamaterials: a gradient-enhanced continuum approach. *Comput. Methods Appl. Mech. Eng.* **452**, (2026).
7. Logg, A., Mardal, K.-A. & Wells, G. *Automated Solution of Differential Equations by the Finite Element Method: The FEniCS Book*. (Springer Science & Business Media, 2012).
8. Balay, S. *et al. PETSc Users Manual*. http://www.mcs.anl.gov/petsc (2006).
