## Supplementary Note 2 for "Cyst-like intraluminal hyaluronic acid accumulation inhibits breast epithelial cell invasion via pressure-driven ECM compaction"

### Supplementary Note 2: Computational Model for Lumen Formation

In this section, we outline a cellular-based computational model for lumen formation. The model extends the traditional two-dimensional vertex model in which the cross-section of cells in a tissue are characterized as deformable polygons with some energetic cost for not conforming to a target shape [1–5]. Moreover, in the traditional approach, cell-cell interfaces are shared so that the tissue is fully confluent, i.e, there are no gaps between cells. We extend this traditional vertex model to allow for gaps between cells such that they no longer share interfaces. We also incorporate a flow network between the cells that can help drive lumen formation. See Fig. S1. With the new extended vertex model, we use experimental data obtained from control and HA over-expressing spheroids to inform our model and refine key parameters based on empirical observations.

#### Extended Vertex Model

Traditional vertex models with small numbers of cells have been used to investigate lumen formation [6]. Given that traditional vertex models have shared cell-cell interfaces, the location of the lumen is pre-determined by explicit gaps. With our extended vertex model, where none of the cells share interfaces, a pipe flow network of paracellular fluid between the cells can now help determine where the lumen form in response to both cellular deformations and fluid accumulation, i.e., the location of lumen are no longer pre-determined. When examining both roles, prior work demonstrates that early epiblast lumen formation is driven by mechanical forces from actin remodeling, while later expansion is due to the osmotic pressure-driven accumulation of fluid [7]. The pressure-driven mechanism is further supported by recent work from Mukenhirn et. al. that demonstrates later lumen expansion is dominated by hydrostatic pressure from ion accumulation [8]. We assumed that the simple pressure term can represent global behavior, as ion transport is governed by both osmotic and hydrostatic pressure [9]. Under this assumption, the pressure differential is a constant across each cell-cell junction in the spheroid, as opposed to modeling a heterogeneous distribution of pressure gradients as a function of location within the spheroid. Additionally, we model cell-cell adhesions as springs that can be ruptured above a critical value of strain. For our two-day lumen formation simulations, we simplified the model by excluding cellular intercalation T1 transitions and cell division [10–12].

##### Energy Functional.

To predict the location and shape of lumen using energy minimization, we begin with one of the standard energy functionals for two-dimensional vertex models for cells indexed by $t$ [4, 5] though allowing for cells to swell as the spheroid takes on fluid:

$$E_{\text{cell}} = \sum_t K_A (A_{t(\text{int})} - a_{t(\text{int})} A_0)^2 + \sum_t K_P (P_{t(\text{int})} - p_{t(\text{int})} P_0)^2$$

$$+\sum_t K_A(A_{t(\text{per})} - a_{t(\text{per})}A_0)^2 + \sum_t K_P(P_{t(\text{per})} - p_{t(\text{per})}P_0)^2. \tag{S1}$$

The area terms account for the bulk deformability of the cells while the perimeter terms account for the contractility of the actomyosin cortex. Moreover, $A_0$ and $P_0$ represent the area and the perimeter of the regular hexagon respectively. Note that we have assumed that the bulk elasticity ($K_A$) and the contractility ($K_P$) are the same for all cells for simplicity. Moreover, we have separated the interior cells of the spheroid, labelled with the subscript *int*, and the cells at the periphery, or boundary, of the spheroid, labelled with the subscript *per*. We do so because experiments demonstrate that the periphery cells do not swell as much as the interior cells. To take into account this swelling as the spheroid accumulates fluid, the dimensionless $a_{t(int)}$, $a_{t(per)}$, $p_{t(int)}$, $p_{t(per)}$ denote the fractional amount of swelling of the areas and perimeters of both interior and periphery cells. Table S1 shows specific values for different phases to be discussed below.

Given that the system is a spheroid with a finite size, we also include an interfacial tension term for cell interfaces at the periphery via [13]

$$E_{\text{spheroid}} = K_I P_{\text{sph}}, \tag{S2}$$

where $P_{\text{sph}}$ denotes the perimeter of the spheroid and $K_I$ the interfacial tension. Finally, as the spheroid takes on more fluid and the cellular deformations become more extreme we add the following contribution to the energy functional,

$$E_R = \sum_i K_R(v_i - v_t)^2 = \sum_i K_R\left((x_i - x_t)^2 + (y_i - y_t)^2\right), \tag{S3}$$

where $v_i$ denotes vertex $i$ with coordinates $(x_i, y_i)$. This energetic contribution insures that the vertices coordinates deform smoothly as the cells expand.

Now that we have described the energy function of the traditional vertex model, we now move to our extended vertex model for lumen formation that includes 1) an osmotic pressure-driven lumen expansion phase [7], and 2) novel determination of lumen location initiation as a function of cell-cell adhesion strength. As inspired by Indana, *et al.*, two phase model, during Phase 1, cell junction stiffness decreases, the distance between cells increases, and lumens begin to form. Our experimental data suggests that the bulky glycocalyx produced by HAS overexpressing cells reduces cell-cell adhesions (Fig. S2). During Phase 1, we assumee the junction springs above a critical length will break, potentially creating lumen reservoirs at those locations [9]. During Phase 2, all remaining junction stiffness values are set to higher values, as the remaining cell-cell junctions stiffen, and lumens begin to grow with increasing hydrostatic pressure. Finally, given our quasistatic approach of energy minimization in which dynamics is modeled by a change in parameters in the energy functional, each Phase is broken down into 4 sub-phases to allow for the encoding of time-scale behavior to simulate the 2 day growth of spheroids cultured *in vitro*.

**Table S1**: Constraints used in the cell energy functional for cell swelling expressed as ratios, where $1.1 \cdot v_0 = (1.1 \cdot x_0, 1.1 \cdot y_0)$ and $v_{a-b}$ represents the vertex coordinate at phase $a - b$.

| **Variable** | $a_{t(\text{int.})}$ | $p_{t(\text{int.})}$ | $a_{t(\text{peri.})}$ | $p_{t(\text{peri.})}$ | $v_t = (x_t, y_t)$ |
|---|---|---|---|---|---|
| Phase 1-1 | $(1.04)^1$ | $\sqrt{(1.04)^1}$ | 1 | 1 | $1.1 \cdot v_0$ |
| Phase 1-2 | $(1.04)^2$ | $\sqrt{(1.04)^2}$ | 1 | 1 | $1.1 \cdot v_{1-1}$ |
| Phase 1-3 | $(1.04)^3$ | $\sqrt{(1.04)^3}$ | 1 | 1 | $1.1 \cdot v_{1-2}$ |
| Phase 1-4 | $(1.04)^4$ | $\sqrt{(1.04)^4}$ | 1 | 1 | $1.1 \cdot v_{1-3}$ |
| Phase 2-1 | $(1.04)^5$ | $\sqrt{(1.04)^5}$ | 1 | 1 | $1.1 \cdot v_{1-4}$ |
| Phase 2-2 | $(1.04)^6$ | $\sqrt{(1.04)^6}$ | 1 | 1 | $1.1 \cdot v_{2-1}$ |
| Phase 2-3 | $(1.04)^7$ | $\sqrt{(1.04)^7}$ | 1 | 1 | $1.1 \cdot v_{2-2}$ |
| Phase 2-4 | $(1.04)^8$ | $\sqrt{(1.04)^8}$ | 1 | 1 | $1.1 \cdot v_{2-3}$ |

| **Coefficient** | **Symbol** | **Value** | **Standard Units** | **Functional Form** |
|---|---|---|---|---|
| Area Modulus | $K_A$ | 0.5 | $\text{Pa}^*/\mu\text{m}^*$ | $K_A(A_i - A_0)^2$ |
| Perimeter Modulus | $K_P$ | 0.5 | $\text{Pa}^* \cdot \mu\text{m}^*$ | $K_P(P_i - P_0)^2$ |
| Resistance Coefficient | $K_R$ | 0.2 | $\text{Pa}^* \cdot \mu\text{m}^*$ | $K_R[(x_i - x_t)^2 + (y_i - y_t)^2]$ |
| Spheroid Surface Tension | $K_I$ | 1 | $\text{Pa}^* \cdot \mu\text{m}^{2*}$ | $K_I P_{\text{sph}}$ |
| Fluid Energy | $K_L$ | 1 | $\mu\text{m}^*$ | $K_L \cdot pA_{\text{pipes}}$ |

**Table S2**: Energy functional parameters for the cells and the paracellular fluid with the exception of the cell-cell adhesion spring strengths. Note that the units are in arbitrary simulation units denoted by asterisk $*$ that will be converted to biophysical units when making direct comparisons with experiments.

| **Stage** | **Junction State** | **Process** |
|---|---|---|
| Phase 1 | Leaky junction phase | Molecule transport through pipe network |
| Phase 2 | Tight junction phase | Selected pipes become lumen and expand |

**Table S3**: Two-phase model of lumen formation

To differentiate between Phases 1 and 2, there energy functional contribution is used in Phase 1 and another for Phase 2:

$$E_{\text{pipes}} = -K_L \cdot pA_{\text{pipes}}, \tag{S4}$$

$$E_{\text{lumen}} = -K_L \cdot p \sum_{lumen} A_{\text{lumen}} \tag{S5}$$

The key difference between these energy functions is that in Phase 1, the area of paracellular fluid between the pipes is represented as ($A_{\text{pipes}}$), while in Phase 2, only selected pipes exceeding a critical length threshold are included in the lumen area calculation ($\sum A_{\text{lumen}}$). Table S2 summarizes energy functions we introduced with their coefficients and relative units.

To complete the model, as the cells no longer share interfaces, we incorporate cell-cell adhesions between cells with springs between the vertex of one neighboring cell to the closest vertex of any other neighboring cell via

$$E_{\text{junction}} = \sum_{i,j \in G(\alpha,\alpha)} \frac{1}{2} K_{S1} d(v_i, v_j)^2 + \sum_{i,j \in G(\alpha,\beta)} \frac{1}{2} K_{S2} d(v_i, v_j)^2 + \sum_{i,j \in G(\beta,\beta)} \frac{1}{2} K_{S3} d(v_i, v_j)^2 + \sum_{i,j \in G(\text{bd},\text{bd})} K_B d(v_i, v_j)^2, \tag{S6}$$

where $d(v_i, v_j)$ denotes the distance between vertices $v_i$ and $v_j$ (Fig. S1).

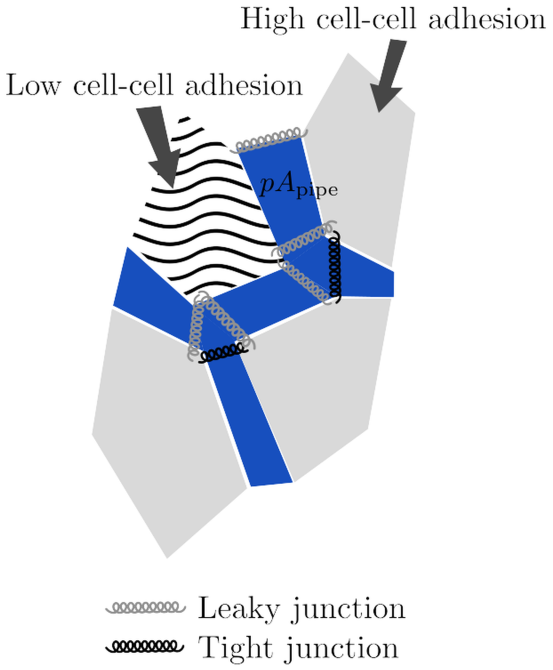


**Fig. S1**: **Cell network model:** Wavy patterned cells represent low cell-cell adhesion, while gray cells indicate high cell-cell adhesion. Junction stiffness corresponds to the stiffness of the springs between cells.

To model the potential for different junctional tensions, $\alpha$ and $\beta$ representing two groups of cells in terms of their cell-cell adhesion strength, and $G(\alpha, \beta)$ denotes junctions connecting cells from group $\alpha$ to group $\beta$. Fig. S1 illustrates the simplest case with two different cell groups. For example, $G(\alpha, \beta)$ corresponds to gray-colored springs with low stiffness, representing leaky junctions during Phase 1. We assumed no hydraulic fracturing/rupturing, thus, osmotic pressure does not break junction but can make junctional tension weaker as captured by different stiffness sets between groups. At the end of Phase 1, stretched springs longer than the set critical junction length are cut and assumed to rupture due to hydrostatic pressure and applied tension to the cell-cell junctions.

#### Differential Cell-cell Adhesion Patterns.

Motivated by previous experiments [14], we implemented three distinct initial spatial cell-cell adhesion patterns for our collagen and suspension models. These previous experiments demonstrated that collagen-embedded cells undergo confinement-induced cellular rearrangement when populations with different cell-cell adhesion properties are combined [14]. These results revealed that confinement within collagen prompts cells within the spheroid to self-organize in which cells with low adhesion strength will preferentially locate at the periphery of the spheroid. While their experiment specifically mixed two different cell types with distinct adhesion properties, we argue below that differential HA production within the spheroid can lead to similar effects.

The biological basis for incorporating intraspheroidal cellular adhesion patterning is supported by several key experimental findings:

1. Overexpression of HAS2 before spheroid formation and the subsequent development of a HA-rich glycocalyx prevents cellular aggregation into spheroids. This suggests a reduction in cell-cell adhesions associated with HAS2 expression (Fig. S2).

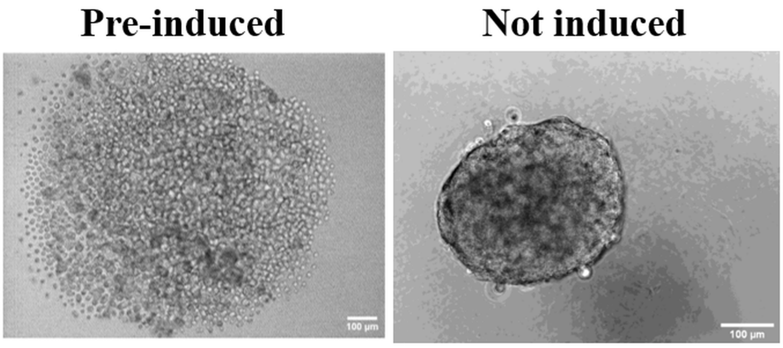


**Fig. S2**: **Experimental image:** Pre-induced cell group does not form spheroid.

2. The HAS2 cell population is heterogeneous in HA production and induced phenotype [15]. Thus we infer that variations in HA production between cells will lead to heterogeneous distribution of cell-cell adhesion strengths. Additionally, heterogeneity in HA production is likely to be amplified in spheroids due to the presence of intraspheroidal metabolic gradients [16] whereby metabolic control of HA production is a key determinate of cell phenotype [15].

3. The Differential Adhesion Hypothesis [17] states that aggregates of cells will self-sort and segregate based on relative adhesion strength. And while self-sorting in spheroid co-cultures can occur in suspension with stronger adhesion in the core when cell proliferation is inhibited [18], work by Cai, et al. [14] indicates that strong matrix confinement promotes high adhesion cells to self-associate at the spheroid core while lower adhesion cells will separate into the outer layer of the spheroid. In the suspension case, it may be that spheroid confinement is enhanced by the interfacial tension of the spheroid, at least for the cell types in Ref. [18].
4. Our experimental data demonstrates that spheroids experience compression due to collagen matrix confinement (main text figure 2). This confinement would induce DAH cell sorting in the matrix embedded conditions. Suspension cultures do not experience matrix confinement and thus cells do not self-sort based on adhesion strength to the extent they do in the collagen-confined spheroids. Moreover, we do not anticipate an enhanced interfacial tension due to the inhibition of cell proliferation.
5. HAS2 overexpression promotes epithelial to mesenchymal transition (EMT) including loss of membrane associated E-cadherin in epithelial cells [19]. There is additional evidence that confinement also promotes EMT and the corresponding loss of E-cadherin [20]. These studies combine to suggest that multiple features present in our experimental system combine to reduce cadherin based cell-cell adhesions and create adhesion strength heterogeneity in our population of cells.

Given these findings, Fig. S3 illustrates the initial cell distribution patterns based on adhesion strength used in our simulation for the two experimental settings—the suspension setting and the collagen setting.

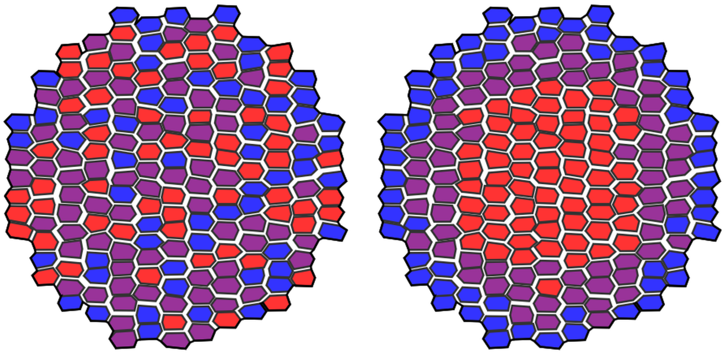

**Fig. S3**: **Initial Cell-Cell Adhesion Heterogeneity Model for spheroids in suspension and for spheroids embedded in collagen:** We assume the spheroids in suspension (left figure) do not experience the same level of effective cell sorting based on adhesion strength that the spheroids embedded in collagen (right figure), while collagen embedded groups do undergo effective cell sorting based on adhesion strength. Blue cells represent low cell-cell adhesion, purple cells indicate intermediate (or medium) adhesion, and red cells represent higher cell-cell adhesion.

| **Parameters** | **Sample Types** | | | |
|---|---|---|---|---|
| | HAS2-induced Suspension | HAS2-induced Collagen | HAS2 non-induced Collagen | Parental (control) |
| $K_S$ of $G(L, L)$ | 0 | 0 | 1 | 2 |
| $K_S$ of $G(L, M)$ | 1 | 1 | 2 | 3 |
| $K_S$ of $G(L, H)$ | 1.5 | N/A | N/A | N/A |
| $K_S$ of $G(M, M)$ | 2 | 2 | 3 | 4 |
| $K_S$ of $G(M, H)$ | 2.5 | 2.5 | 4 | 5 |
| $K_S$ of $G(H, H)$ | 3 | 3 | 5 | 6 |
| $K_S$ at Phase 2 | 10 | 10 | 10 | 10 |
| Critical junction length | 0.5 | 0.5 | 0.5 | 0.5 |
| $p$ | 0.001 | 0.001 | 0.001 | 0/0.001 |

**Table S4**: Junction stiffness ($Pa^* \cdot \mu m^*$), critical junction length ($\mu m^*$), and osmotic pressure ($Pa^*$) for each sample type in arbitrary simulation units (scaled relative to experimental values).

In our simulations, we used the pattern shown in Fig. S3 (right) as the initial configuration for collagen spheroids during Phase 1. We also tested alternative patterns that transitioned between the left and right configurations in collagen spheroids during the subphases of Phase 1 to model a gradual process of confinement-induced cell sorting. Interestingly, we did not observe significant differences in lumen location between an initial sorted population and one that was sorted gradually during phase 1. The consistency in lumen location across the pattern variations, suggests that only the adhesion pattern during the later stages immediately proceeding junction breakage (at Phase 1-3/1-4 in Table S1) is crucial for determining final lumen morphology.

#### Differences in model parameters for different experimental environments.

We assume: 1) HAS2 influences junction stiffness and promotes cell detachment, and 2) HAS2 promotes the production of HA that elevates osmotic pressure on the interior of the spheroid which in turn drives fluid flow along the osmotic gradient creating a difference in fluid pressure that forms lumens. These assumptions are supported by experimental evidence. For the first assumption, our experiments showed that the pre-induced cell group does not form spheroids, as demonstrated in Fig. S2. For the second assumption, HA is well reported to create an increase in osmotic pressure and subsequently generate hydrostatic pressure to deform the surrounding tissues [21] [22]. Additionally, published measurements [8, 23] and our own experiments (Main Text Figure 3) demonstrate elevated hydrostatic pressure within mature lumens. Using these assumptions, we applied different parameters for the different experimental environments as shown in Table S4:

#### Numerical estimation of the pressure.

We applied the following scale conversions to convert to experimental units given the arbitrary simulation units [8, 23]. We do so because our simulation outputs require calibration to experimental measurements for comparison [24]. We start with the pressure difference between the two ends of pipe of radius $r$ and length $l$, or $p$ (Pressure) $= \frac{8\mu Q l}{\pi r^4}$, where $\mu$ denotes viscosity and $Q$ denotes the volumeric flow rate. Then,

$$\text{Scale factor} = \frac{16\,\mu\text{m}^*}{200\,\mu\text{m}} = \frac{1.0}{12.5}\,\mu\text{m}^*/\mu\text{m}, \tag{S7}$$

$$r = 1.0\,\mu\text{m} \times \frac{1.0}{12.5} = 0.08\,\mu\text{m}^*, \tag{S8}$$

$$l = 3.2\,\mu\text{m} \times \frac{3.2}{12.5} = 0.256\,\mu\text{m}^*, \tag{S9}$$

$$Q = 900\,\mu\text{m}^3/\text{s} \times \left(\frac{1}{12.5}\right)^3 = 0.4608\,\mu\text{m}^{3*}/\text{s}, \tag{S10}$$

$$\text{Viscosity}(\mu) = 5.1 \times 10^{-9}\,\text{kg}/(\mu\text{m}\cdot\text{s}) \times 25 \times \frac{16}{8} = 1.03 \times 10^{-7}\,\text{kg}/(\mu\text{m}^*\cdot\text{s}), \tag{S11}$$

$$p\ (\text{Pressure}) = 35\,\text{Pa} \approx 7.554 \times 10^{-4}\,\text{Pa}^* \approx 0.001\,\text{Pa}^*\ (\sim 40\,\text{Pa}). \tag{S12}$$

Previous *in vitro* systems have found luminal pressure to be ($\sim$ 3 Pa) [23] in pancreatic organoids and ($\sim$ 50 Pa) in MDCKII cysts [8]. Although this calculated pressure value is higher than experimentally measured pressure in the HAS2 spheroids($\sim$ 10 Pa), we are in a physiologically relevant range for in vitro organoids and our sensitivity analysis showed that the parameter $p$ does not significantly affect simulation outcomes even when quadrupled relative to junctional tension. This is due to the strength of energy function differences small enough amount for osmotic pressure $< 1$ compared to spring network ($< 100$ or more depending on the stiffness) in the simulation. Also, we would like to note that fluid flows into leaky junctions without affecting junction stiffness or weakening them through excessive water flow, thus in the simulation the pressure term plays a more passive role rather than an active one. Furthermore, while we assumed a spheroid diameter of approximately 200 $\mu m$ with 200 cells in our baseline calculations, experimental spheroids often reached 300 $\mu$m or larger with 10000 cells, suggesting that our selected value of $p = 0.001$ remains physiologically appropriate within the range for the system being modeled. Similar results shown in the paper [8] ZO-KO group does not develop large lumen due to high stiffness which could be related to the collagen sample obtained from HAS2-overexpressed with no induction.

#### Other simulation details.

As mentioned previously, we used a vertex tethering/resistance function (Equation S3) to help regulate the positions of the vertices as the spheroids evolve. As shown in Table S1, the swelling ratio is set to 1.1. This value yields a cumulative expansion of approximately 2.14 ($(1.1)^8 \approx 2.14$), which is greater than the 87.7% expansion observed in suspension Fig. S4. We chose this higher value to ensure sufficient expansion of both the pipe network and cells, creating adequate space for lumen formation. This function can be interpreted as uniform cell displacement resulting from hydrostatic pressure between the spheroid center and its exterior. Our sensitivity tests with slightly different values confirmed that this parameter does not affect the location of lumen formation.

For our simulation, we incorporated junction stiffness values ($\frac{1}{2}K_S$) within a range of 0-5 Pa* (Table S4), based on experimental observations from tight junction studies [8]. We derived these relative stiffness values from laser ablation experiments that measured vertex recoil velocities: approximately 1.5 $\mu$m/s for wild-type, 3.5 $\mu$m/s for claudin knockout (2.3× WT), and 10.5 $\mu$m/s for zonula occludens knockout (7.0× WT) conditions. Since higher recoil velocities indicate weaker junctions (increased paracellular leakage), we established an inverse relationship where the experimental velocity ratios map to junction stiffness values: wild-type conditions correspond to tight junction stiffness ( 5.0 Pa*), claudin knockout to normal junction function ( 2.2 Pa*), and zonula occludens knockout to weak junctions ( 0.7 Pa*) that promote lumen formation.

To set the absolute scale, we used established cell-cell junction mechanics: forces of ∼100–120 nN across contact areas of ∼50 $\mu$m$^2$ yield junction stresses of approximately 2–2.4 kPa [25, 26]. Converting to our 2D simulation framework, where forces act along spring connections rather than across 3D contact areas, these stresses correspond to approximately 1.25–1.5 Pa* in our simulation units (accounting for the geometric conversion factor of 25 to convert from 3D area-distributed to 2D line-concentrated forces). Our simulation parameter range captures the biological spectrum: 0 Pa* represents completely disrupted junctions with maximal leakage, 1–2 Pa* corresponds to normal junction function (consistent with literature values), and 3–5 Pa* represents tight junction states (WT-like) during barrier formation.

We note that our simulation scale (200 cells, ∼1 $\mu$m$^{2*}$ average cell area) differs from typical experiments (∼10,000 cells, ∼50 $\mu$m$^2$ average cell area). However, our approach using relative stiffness values allows us to model the key factors determining lumen size and location while maintaining computational efficiency.

##### Factors determining lumen size and location.

Based on our model and tests using different parameters under the same energy functions, we found that the lumen size is determined by both junctional tension and critical length. The lumen location is influenced by the leaky cell pattern that emerges near the end of Phase 1. Also, the size of the lumen is related to critical length which explains how much neighboring cells can be close each other when they are far apart. Given the experimental determination of osmotic pressure, we leave the study of different osmotic pressure for future work.

##### Determining the cell shape index from experimental images.

To assign appropriate values, we measured cell area ratios from Day 0 to Day 2 using two-dimensional cross-section images. We observed that parental samples exhibited area shrinkage, which was also reflected in the overall spheroid area, indicating that no cell swelling occurred and the gaps between cells decreased. This phenomenon can also be observed in Fig. S4. In contrast, both spheroids embedded in collagen and spheroids in suspension samples showed significant swelling in both inner spheroid cells and outer (near the periphery) cells. Given the variation in cell swelling, we made a simplifying assumption by selecting a scaling factor of $(1.04)^8 \approx 1.37$ for inner cells while assuming no change in area for outer cells. This approximation for inner cells aligns with the measured values of 1.38 shown in Table S5. The periphery cells remained at the same target area since collagen samples decrease in area while suspension samples increase in area as compared to the Parental Day 0 sample. While we could have applied different values for each sample and even each cell, we deliberately used a uniform value across all samples so that heterogeneous cell swelling as a critical mechanism in determining lumen location is left for subsequent studies.

We fixed the ratio of perimeter/$\sqrt{\text{area}}$ (shape index) as shown in Table S1. This decision was based on shape index measurements in Table S6, which demonstrated a consistent shape index of approximately 4.5 across all groups. However, we excluded the Day 2 collagen embedded HAS2 samples from this analysis, as we hypothesized potential interactions between collagen fibers and the spheroid boundary which could be additional factors not accounted for in our basic simulation model.

We used convex polygons to model each cell; however, a shape index of 4.5 corresponds to that of a triangle as a regular polyhedron. Rather than using these raw values, we quantified cell morphology using contour analysis. We derived a representative cell shape by averaging observed cell morphologies and fitting them to a hexagon, as illustrated in Fig. S5. This representative shape has a shape index of 3.85. We fixed this number by applying the same ratio of 1.04 for area and $\sqrt{1.04}$ for perimeter relative to the regular hexagon. However, the average shape index is far from the hexagon due to effects from the other energy contributions.

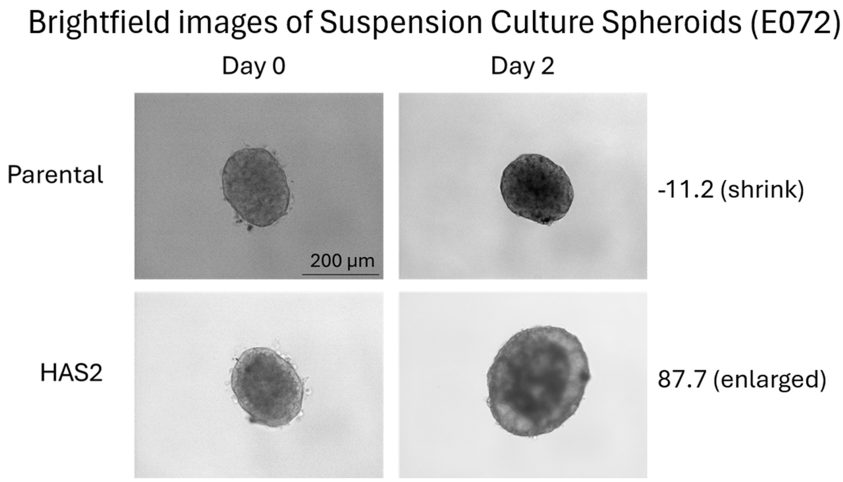


**Fig. S4**: **Experimental image:** Brightfield images of Day 0 and Day 2 size.

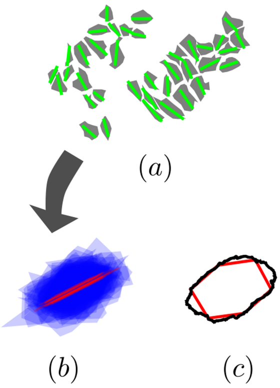


**Fig. S5**: **Representative Cell Shape Measurement:** (a) To simulate cells that more closely resemble experimental conditions, we selected interior cells from the parental sample, aligned them along their axes (b), and overlapped them. We then applied a 65% contour threshold and fitted the resulting shape to a hexagon (c).

**Table S5**: Cell area ratios relative to Day 0 samples.

| Condition | Inner Cells | Outer Cells |
|---|---|---|
| Collagen Parental | 0.81 | 1.04 |
| Suspension Parental | 0.91 | 1.23 |
| Collagen HAS2 | 1.38 | 0.78 |
| Suspension HAS2 | 1.70 | 1.49 |

Values calculated as (Sum of sample mean areas)/(Number of samples × Day 0 reference area) from Table S6.

### Code Availability

The Mathematica notebooks used to generate the vertex-model simulations in Fig. 4(c) and Fig. 10(a),(b) of the main text are available in Zenodo at https://doi.org/10.5281/zenodo.24411408.

### Data Availability

The simulation data (lattice configuration a nd s imulation r esults f or F ig. 4 (c) and Fig. 10(a),(b) of the main text) are available in Zenodo at https://doi.org/10.5281/zenodo.24411408.

**Table S6**: Morphological Measurements Across Culture Conditions and Sample Types

| Condition | Sample | Region | Area ($\mu m^2$) | Peri. ($\mu m$) | Shape Index |
|---|---|---|---|---|---|
| **Day 0 Collagen Embedded** | NeOR | Inner | 54.8 | 31.6 | 4.37 |
| | | Outer | 71.8 | 38.3 | 4.63 |
| | HAS2 | Inner | 43.0 | 29.3 | 4.57 |
| | | Outer | 53.1 | 32.1 | 4.71 |
| **Day 2 Collagen Embedded Parental** | NeOR | Inner | 44.3 | 28.8 | 4.38 |
| | | Outer | 74.6 | 37.4 | 4.52 |
| | NeOR* | Inner | 25.4 | 20.5 | 4.20 |
| | | Outer | 27.8 | 23.6 | 4.66 |
| **Day 2 Collagen Embedded HAS2** | HAS2 | Inner | 59.5 | 33.5 | 4.39 |
| | | Outer | 52.8 | 33.7 | 4.73 |
| | | Lumen | 65.8 | 39.0 | 4.89 |
| | HAS2 | Inner | 59.6 | 34.7 | 4.62 |
| | | Outer | 30.2 | 29.4 | 5.52 |
| | | Lumen | 64.9 | 44.1 | 5.60 |
| **Day 2 Suspension Culture Parental** | NeOR | Inner | 49.4 | 29.2 | 4.26 |
| | | Outer | 69.8 | 39.9 | 4.99 |
| | NeOR | Inner | 50.5 | 29.1 | 4.22 |
| | | Outer | 106.1 | 42.9 | 4.34 |
| **Day 2 Suspension Culture HAS2** | HAS2 | Inner | 71.0 | 37.5 | 4.49 |
| | | Outer | 64.5 | 35.4 | 4.49 |
| | | Lumen | 62.2 | 32.8 | 4.28 |
| | HAS2 | Inner | 72.3 | 36.5 | 4.41 |
| | | Outer | 91.3 | 38.9 | 4.22 |
| | | Lumen | 50.7 | 30.8 | 4.46 |
| | HAS2 | Inner | 76.5 | 36.1 | 4.23 |
| | | Outer | 82.5 | 38.5 | 4.32 |
| | | Lumen | 67.3 | 37.0 | 4.66 |

*Not included in the average data in Table S5 because the area is almost half that of Day 0. The mean data were calculated for each group.
